# GAI MoRFs Regulate Cleft and Channel Binding Pathways for Gibberellin in GID1A

**DOI:** 10.1101/2020.12.15.422840

**Authors:** John Patterson, Charles C. David, Marion Wood, Xiaolin Sun, Donald J. Jacobs, Erik H. A. Rikkerink

## Abstract

**Abstract:** The hormone gibberellin (GA) promotes arabidopsis growth by enhancing binding between GA Insensitive DELLA transcriptional repressors and GA Insensitive Dwarf 1 (GID1) receptors to regulate DELLA degradation. The binding mechanism for GA was elucidated by employing a computational study of dissociations of the N-terminus of the DELLA family member GAI (GA Insensitive transcriptional repressor) from the GID1A receptor in the presence and absence of bound GA, and of GA from GID1A in the presence and absence of GAI. The tRAMD method was employed to deduce egression pathways for a diverse set of GA molecules (GA^(x)^). Two pathways in the form of a newly identified cleft and a previously identified channel are prevalent. The cleft pathway is open in the absence of GAI. Upon GAI binding, the cleft route is blocked, resulting in a slower process for GA^(x)^ to exit and enter the binding pocket through the channel. Several binding pocket residues are identified as gate-keepers to the channel. Molecular recognition features (MoRFs) found in the disordered signaling protein GAI affect GA^(x)^ binding and GID1A dynamics. A three-step synergistic binding cycle is proposed where GAI MoRFs regulate the process. Rapid binding takes place through the cleft where little to no distinctions are made between major and less active forms of GA^(x)^. After GAI is bound to the GA^(x)^ · GID1A complex, the channel supports a rectification process that increases the retention of major active forms of GA within the binding pocket. Both the cleft and channel contact residues to GA^(x)^ are markedly conserved in a GID1 phylogeny, suggesting this binding process in the GID1 · DELLA GA-receptor complex represents a general paradigm for GA binding. Non-specific GA binding assists binding of GAI, which then helps to select the major active forms of the hormone and induce a downstream signalling cascade in response to bioactive GA.

**Non-expert Summary Statement:** Gibberellins are plant hormones essential for growth and development. The DELLA proteins are a disordered family of repressors that transcriptionally repress GA responsive genes. Degradation of DELLA proteins in response to GA results in GA-responsive genes being upregulated. Binding of GA to the GA-Insensitive Dwarf 1 receptor (GID1) facilitates binding of DELLA to the GA · GID1 complex. Through computational modelling and phylogenetic analyses, we identified a new GA binding cleft that is blocked by DELLA binding and a three-step mechanism for the GA · DELLA · GID1 complex that also involves the known GA binding channel. We propose a dual (cleft/channel) pathway that allows access to the binding pocket as a paradigm for selection of specific GA forms among a mixture of major active and inactive forms. The cleft is less selective, but preference for active GA in the binding pocket of GID1A is amplified by expunging inactive GA forms, followed by recruiting active forms through the more selective channel. This mechanism allows plants to sense concentration changes of GA with high specificity to enable certain GA variants to trigger specific signalling events. These novel insights into the receptor mechanism in part may explain the large number of different GA forms that exist in nature.

**Graphical Abstract:** 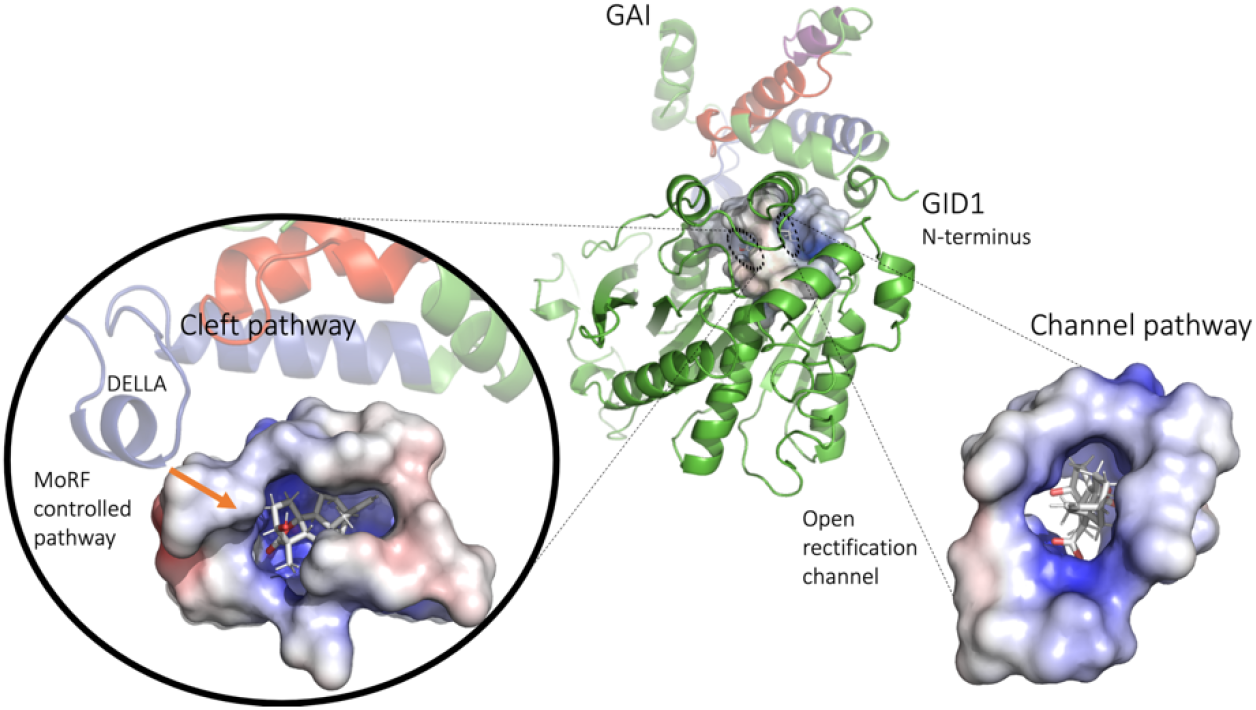

## Introduction

It is well known that protein function does not only arise from highly structured parts of proteins but also depends on regions that have flexibility (1–3). Furthermore, there is now general recognition that highly flexible intrinsically disordered regions found in many proteins can be critical for function. Disordered regions typically contain short semi-rigid segments that appear to play key roles in establishing and controlling interactions with protein partners. Commonly known as <u>Mo</u>lecular <u>R</u>ecognition <u>F</u>eatures (MoRFs), these motifs often show promiscuous folding and binding activities that are enabled by their environments (4, 5). The large number of accessible conformations in disordered proteins allows for binding promiscuity and/or a fine gradient of responses. The DELLA protein GAI, one of 5 members in Arabidopsis, was first identified through its function as a growth-repressing regulatory protein that operates through the gibberellin response pathway (6). Subsequently the DELLA family was identified as a key site of mutations in several grass species that have led to the so called ‘Green Revolution’ of high-yielding dwarfed varieties. The dwarf phenotype stems from interruption of the interaction between the Gibberellin (GA) receptor GA-Insensitive Dwarf1 (GID1) and a DELLA protein (7). Over the next two decades the DELLA family was shown to interact with a large group of protein cohorts involved in a myriad of processes such as stress response (8), plant defence (9–11), jasmonate signalling (12, 13), light perception mechanisms (14, 15), photosynthesis (16), stomatal conductance (17), circadian clock mechanisms (18), nitrogen assimilation (19), iron uptake (20) and protein trafficking (21). The range of signalling pathways interacting with DELLA proteins now covers virtually the entire suite of plant hormones (22, 23), earning them the title of “master regulators of growth and development” (24).

The diversity of roles carried by this relatively small plant protein family is predicated upon its structural plasticity and two main components. The structurally more rigid C-terminal GRAS (Gibberellic-Acid Insensitive, Repressor of GAI and Scarecrow) domain is able to bind to many transcriptional regulatory partners. The intrinsically disordered N-terminal domain contains the family’s eponymous conserved DELLA motif. We have previously proposed that the disordered nature of the N-terminal domain, along with three short conserved motifs that act as MoRFs, are critical to promote its hub function capabilities (4, 5, 25). While these MoRF sites are known, their intrinsic role in molecular recognition is incompletely characterized. Elucidating the molecular details that describe how the DELLA protein interacts with the GID1 protein family will help us understand how this protein complex can play such a key role in plant food production and facilitate any future manipulation of this complex.

A GID1A centric hypothesis for the GA binding mechanism was initially developed based on the GA · GAI · GID1A complex (gDG-where g refers to the hormone GA and D to the DELLA protein GAI) that was crystalized (PDB-code: 2ZSH). This proposed mechanism postulated a large-scale motion in the flexible N-terminal region of GID1A that was thought to act as a lid that opens and closes onto the (previously) enzymatic core of GID1A (26, 27). Upon GA binding to this open lid state of GID1A, the lid was proposed to subsequently close to form a cap. This capping was deemed a necessary step to allow the GAI (DELLA) protein to bind, while re-enforcing interactions between GA and the N-terminal extension of GID1A to maintain a stable gDG complex. This sequence of events involving the intrinsic conformational motions of GID1A and GA binding was proposed to create a recognition site for GAI. Once the GAI protein is bound to the GA · GID1A complex to form gDG, it attracts a key component of the polyubiquitination system and is subsequently degraded by the ubiquitin-mediated proteasomal process. A consequence of this proposed mechanism is that once GAI is bound, the capped GID1 lid remains closed, preventing GA from leaving the binding pocket until the GAI protein is no longer attached, typically as a result of proteolytic degradation.

The proposed capping of the GID1 N-terminal lid by DELLA proteins was challenged by Hao *et al*. (28) through molecular modelling of the gDG complex. Without the GID1 “lid” opening up, it was shown that there is sufficient space for GA to enter and exit the binding pocket through a channel that is present in the GAI · GID1A complex. The free energy of binding through this channel was calculated, and key contact residues within the channel were identified. These results imply that an apo DELLA · GID1 complex (aDG) can exist within the cellular environment. Interestingly, Yamamoto et al. (29) experimentally found that by suitable site directed mutations to GID1, GA independent binding of DELLA proteins to the rice GID1 is possible, albeit binding is still stronger in the presence of GA. They also identified weak binding of arabidopsis GID1B and GAI in the absence of GA. Although there is no direct evidence that GA independent DELLA · GID1 binding plays a regulatory role in plant biology, our modelling and analysis suggests that the aDG complex plays an important function as a kinetic intermediate to ensure that the gDG complex contains a bioactive GA.

There is considerable interest in plant biology to better understand the roles of more than 100 variants of GA found in nature (30). The ability of different forms of GA to be converted to other forms through a series of differentially regulated enzymatic steps, including both activation and inactivation steps, makes it difficult to conclusively link bioactivity to the different forms of GA without mutants in key enzymatic steps. For example, is it that some GA variants are transient intermediate states or do they add to a spectrum of biological activity in their own right? The large number of GA variants suggests the possibility of variable specificity for the receptor and/or receptor complex. This high diversity motivated this study to consider distinctly different GA^(x)^ on the binding mechanisms of DELLA to GID1A and GA^(x)^ to GID1A and the potential implications of these steps to arrive at the gDG complex. A fundamental question is how different GA variants bind to GID1 and comparatively, how effective they are in promoting the gDG complex stability required for downstream proteolysis processes.

To glean insight into the role that GA^(x)^ ligand variants play, we investigated the GA^(x)^ · GAI · GID1A complex using the tRAMD computational method (31). Starting with a bound GA^(x)^ in the binding pocket, this accelerated molecular dynamics approach was used to discover egression pathways that enable GA^(x)^ dissociation. In addition to comparing different GA^(x)^ molecules, the role of the DELLA protein MoRFs in the binding of GAI to GID1A in the presence and absence of GA^(x)^ is assessed. Moreover, the dissociation of GA^(x)^ in the absence of GAI is investigated and assessed to deduce the binding cycle that leads to the GA^(x)^ · GAI · GID1A complex.

All modelling in this work considers the GID1A conformation with a closed lid. This closed state is supported by the highly unfavourable free energy profile of the open “lid” (28) and the fact that this is the only conformation with physical evidence from crystalisation. A prominent GA^(x)^ binding pathway through an open cleft in GID1A when GAI is absent is uncovered. The roles of the MoRFs, residues that correspond to conserved motifs which are deleted in dwarf mutants of the DELLA partners, are identified with respect to these pathways. Conceptualized in Figure 1, a complete kinetic cycle is proposed that provides a mechanism for preferential binding of major bioactive GA in the receptor complex. This model provides significant new insights that explain how specific GA regulation can occur within the presence of a large number of GA variants that exist in nature. These insights will also inform strategies to manipulate the protein complex to increase yield without negatively affecting the multitude of plant phenotypes linked to the promiscuous DELLA family.

**Figure 1:**
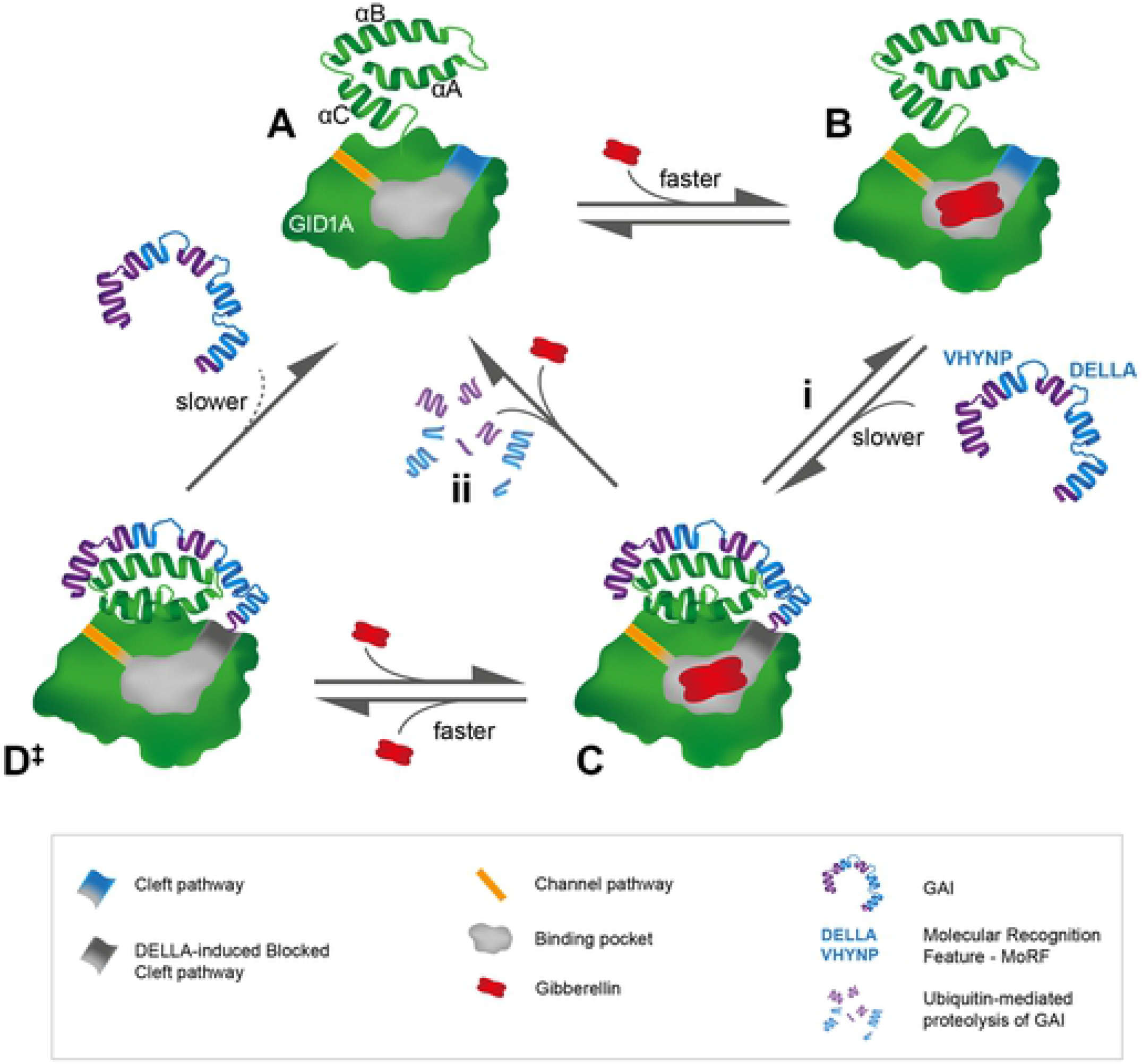
Schematic of the computationally deduced GA binding process. (A) The GID1A receptor has two major routes of GA binding, a readily accessible newly identified cleft (blue) and a long channel (yellow). The GID1A cap (α-helices A to C) is instrumental in defining the binding pocket and it exhibits small-scale motions during the GA binding process. (B) After moving through the cleft or channel, GA binds within the binding pocket in absence of GAI. (C). Upon binding of GAI, the DELLA MoRF covers the cleft access route for GA entry. The GA · DELLA · GID1A complex provides opportunity for degradation of GAI to occur (ii). This can also result in a bound GA · GID1A (i). (D) Due to large differences in time scales, the DELLA · GID1A complex facilitates GA binding through the channel before GAI dissociation occurs. D‡ indicates an unstable intermediate.

## Results

### GA variants considered

Among more than 100 different GA forms found in plants, the dichotomy of active and inactive variants introduced early on (32) is likely too simplistic given the difficulty of tracking which GA variant creates the signalling cascade after a particular form of GA is supplied exogenously. To cover the spectrum of activities that may be inherent in this binary classification scheme, we selected a diverse set of GA^(x)^ that includes GA variants widely referred to as bioactive {GA^(a)^} and as inactive {GA^(i)^}. Additional types of GA are also considered to increase the diversity in molecular moieties. Known GA^(a)^ GA_1_, GA_4_, GA_3_, GA_7_, and recently reported GA_12-16ox_ (33) were chosen to assess active GA containing systems. We also analysed a set of putative inactive forms (GA^(i)^) that represent variants with no evidence of biological activity. Most of these were chosen as a progressive step from GA_12_, the main precursor to all GA, to the various GA^(a)^ examined. In particular, GA_4MeO_, GA_4-16/17ox_, and GA_34_ are all oxidation products of GA^(a)^ and are of interest to understand the signalling control in this system.

### GA binding pocket dynamics during simulation

All production runs for each gDG system retained the bound GA^(x)^ ligand within the binding pocket. Counts of the recurrent hydrogen bonds (H-bonds) and hydrophobic contacts between GA variants and the GID1A residues were monitored. Table 1 summarizes the binding pocking interactions to GA^(x)^ that are largely in common across all the GA variants investigated here. Highlighting these pocket residues outlines the proximity to key parts of GA_3_. TYR247 (see methods for residue numbering) interacts with GA^(x)^ C3-OH, SER116 and GLY115 wrap around the C7 carboxylate. TYR31 has interactions both with OH groups in those GA^(x)^ that have this moiety around C13 and interactions with all GA vicinal C16-C17 bonds, appearing to be aided by pi-pi interactions. This interaction aids in aligning GA inside the GID1A pocket and is a shared moiety across most GA.

**Table 1:**
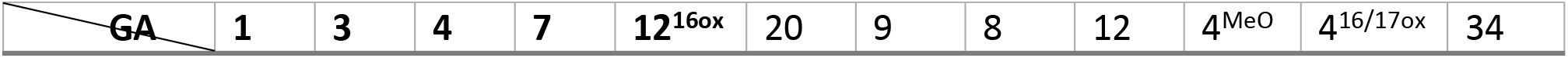
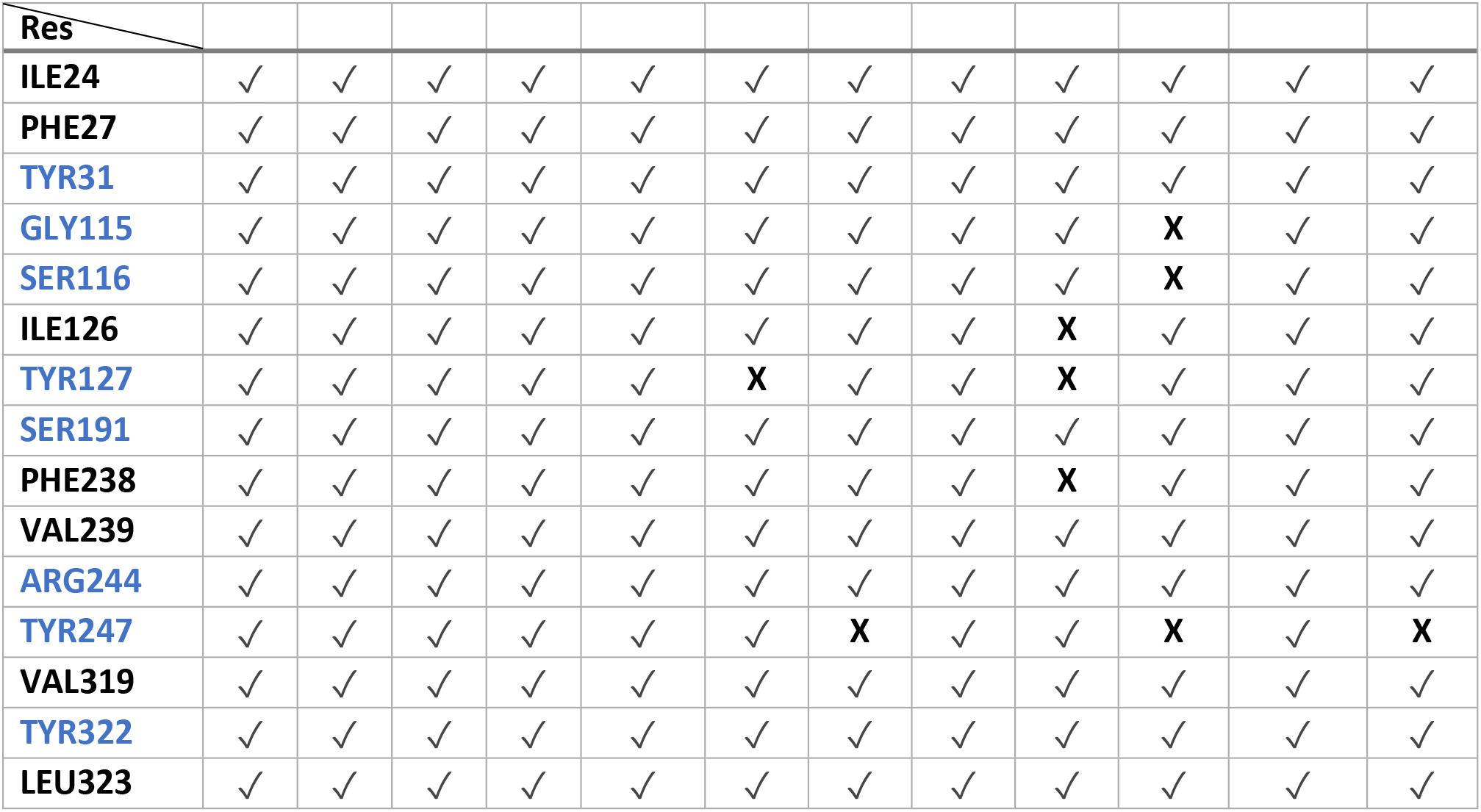
Binding pocket recurrent hydrogen bonds, in blue, and hydrophobic contact residues, in black, from production runs of gDG systems. GA variants in bold are reported bioactive. Residue set generated from GA(a) subset, **X** denotes absence of interaction in given system, while checks denote presence of interaction.

In the GA^(x)^ · GID1 (gG) systems a subset of common interactions was found. With no additional interactions identified, the gG system maintains the same set of interactions except for interactions with TYR31 and TYR247. The loss of the TYR31 and TYR247 interactions enable there to be a less compact environment among the surrounding residues in gG systems compared to gDG systems. Interestingly, the contact residues shown here are highly conserved across the phylogeny of GID1 across diverse plant families (Tables S3–5) suggesting these residues have similar significance beyond the Arabidopsis GAI · GID1A example that we analyze herein.

Conformation changes during the production runs were quantified using the root mean square displacement (RMSD) on carbon alpha atom positions from structurally aligned frames for DELLA and GID1A. The RMSD in the GID1A receptor for all simulations did not exceed 2.5 Å. The RMSD of the disordered DELLA protein was found to have nearly double the RMSD of GID1A, with maximums at 8 Å and an average of approximately 4.5 Å. Breaking this down into the MoRF (4) and inter-MoRF subunits, the DELLA containing MoRF 1 (residues 24 to 55), and VHYNP containing MoRF 2 (residues 69 to 93) have similar levels of RMSD of 1.5 Å. MoRF 3, conserved YL(R/K)LIP residues, (residues 97 to 104) located at the C-terminus end of the GAI N-terminus fragment modelled here has a slightly higher RMSD of 2 Å, which reflects having fewer intra chain interactions compared to the other two MoRFs. As expected, the highest degree of mobility is in the regions between the MoRFs –particularly at the N- and C-terminal regions of the analysed fragment. The relatively lower RMSD found in the MoRFs compared to the linkers are also apparent in the apo form (aDG), meaning GA is removed from the binding pocket.

As expected, the overall degree of mobility in the DELLA protein is reduced in the GA bound forms compared to the apo form. Interestingly, the root mean square fluctuation (RMSF) for the carbon alpha atoms in the region between MoRF 2 and MoRF 3 shows greater mobility with the GA^(a)^ subset when compared with other GA variants analysed, shown in Figure 2 and Figure S1. The GID1A residues were similarly analysed (Figure S9) and were found to be similar between aDG and GA^(a)^ · DG systems. However, the gG systems exhibit an overall increase in RMSF presumably due to the missing interactions with DELLA protein. The regions near pocket residues of GID1A from Table 1 are found to be slightly less mobile in the gDG and gG systems compared to the aDG system.

**Figure 2:**
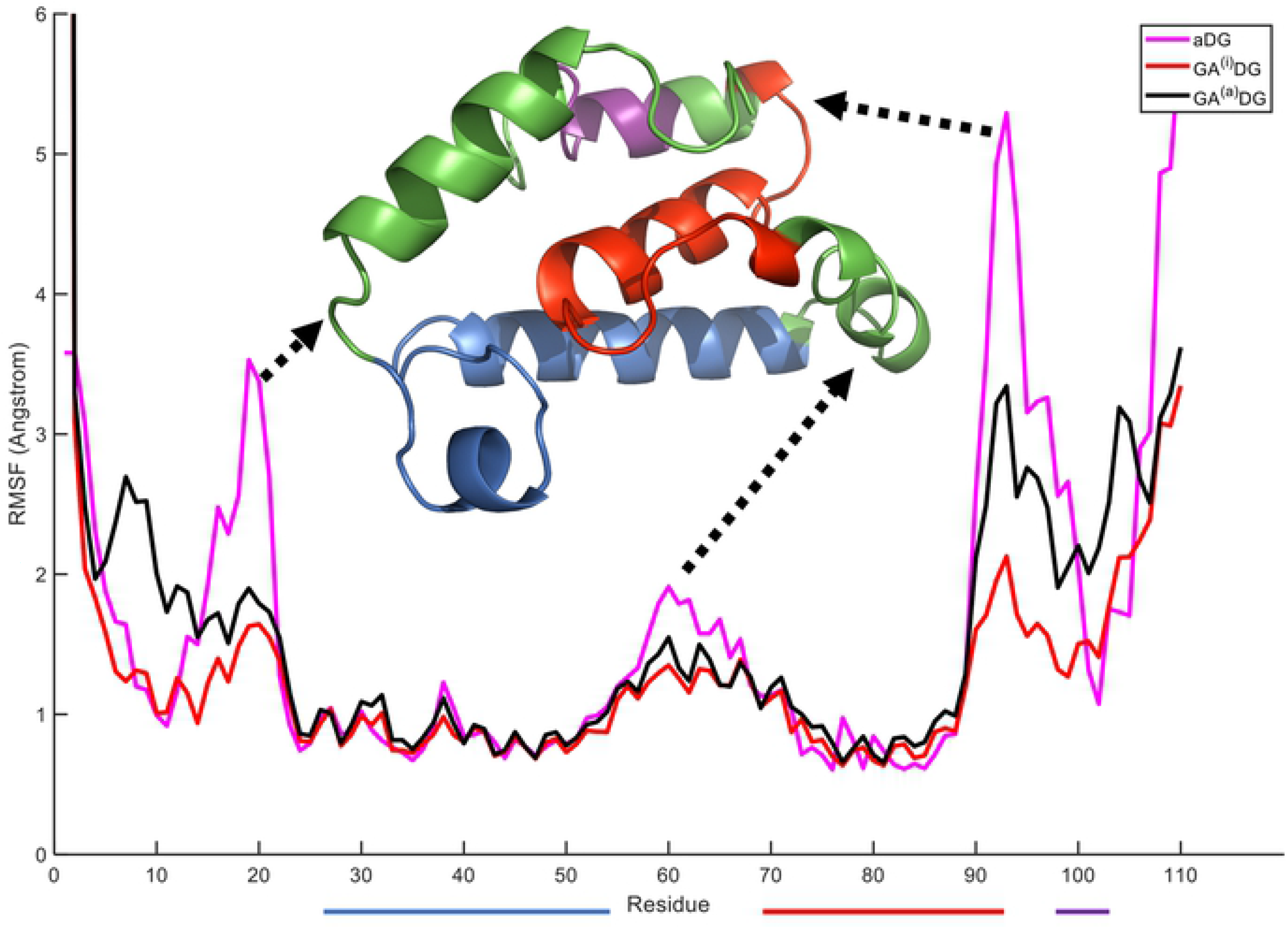
Mean RMSF of production run trajectories in APO GAI · GID1A (magenta) and GA-occupied GAI · GID1A systems. Comparing the subgroups GA^(a)^ (black) with GA^(i)^ (red) containing systems by taking mean RMSF over the duration of the production runs per residue and averaging across all GA within the respective subgroups. The three coloured bars below the x-axis represent the MoRFs and match the colouring of these MoRFs on the GAI ribbon model of the N-terminus above. Dotted arrows match key flexible parts of the model with the mean RMSF values.

### GAx tRAMD analysis: Channel and cleft pathways

After 20 *ns* production runs (computational simulations) were completed, this information was used in tRAMD to identify the GA^(x)^ egression pathways during the dissociation process. We observed that there are essentially two egression pathways, as shown in Figure 3. The longer pathway uncovers the previously reported channel pathway by Hao *et al.* (28). The shorter pathway is dubbed cleft because the opening is readily closed by the presence of DELLA motif and the egressions yielded from this route have fewer interactions overall than in the channel pathway in the GID1A systems.

**Figure 3:**
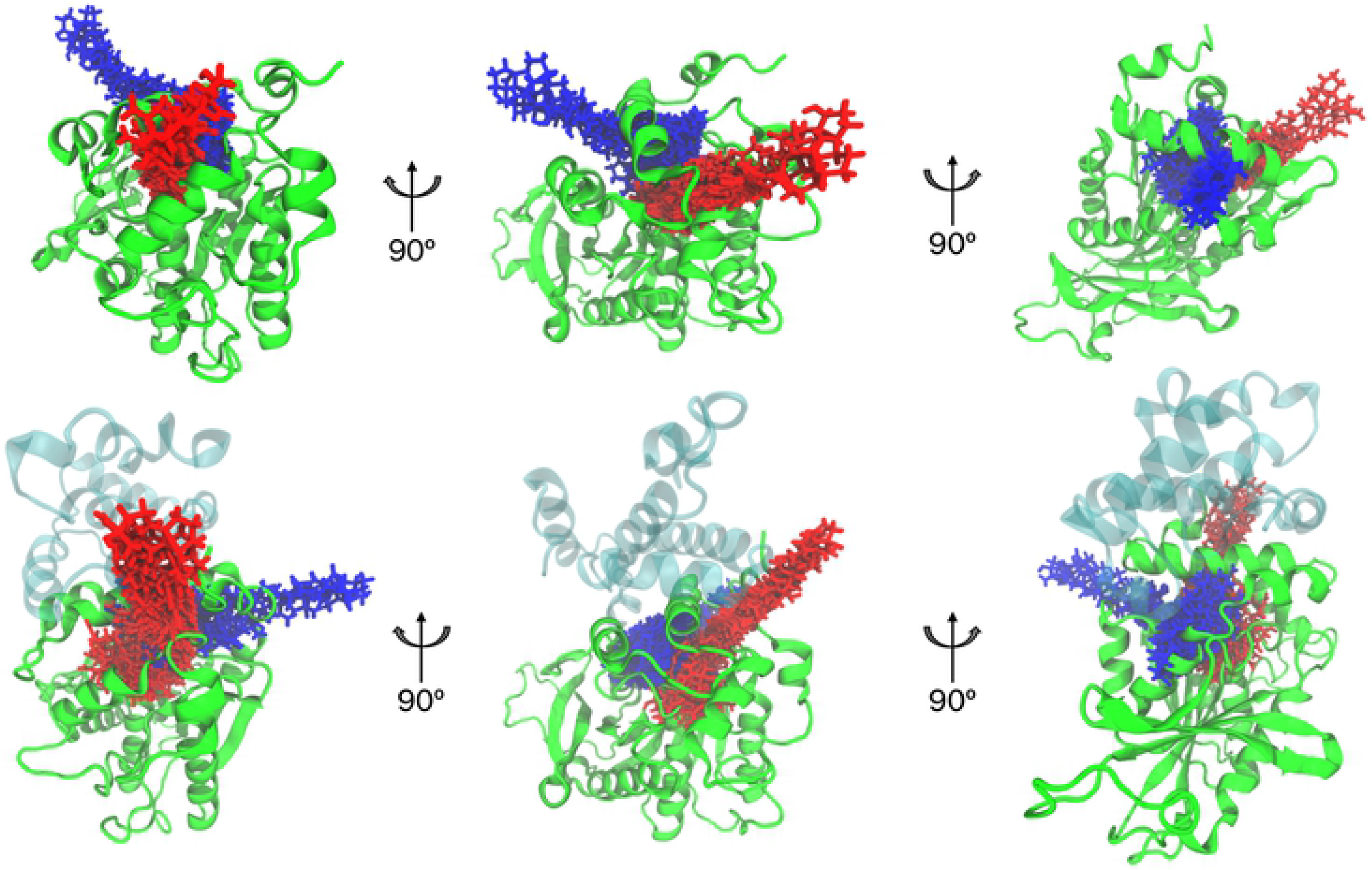
Representations of pathways in GID1A system. Bottom DELLA in cyan and GID1A in green, top GID1A. The blue pathway is a newly discovered GID1 cleft, red is known channel pathway. Middle perspective looks down alpha helix B of GID1A with N-terminal to the right.

It is worth noting that Hao *et al.* used an incomplete gDG structural model, taken directly from the crystal structure (PDB accession code: 2ZSH). Nevertheless, the channel can be identified within that static structure. Subsequently, Hao *et al.* performed free energy calculations using umbrella sampling for GA_4_ as it moved into the binding pocket along this identified pathway. Working inward, from the outside surface of GID1A toward the internal binding pocket, Hao *et. al.* determined the initial residues that interact with GA_4_ along this pathway (that we now call the channel) to be ASP243 and THR240 (note we have not retained all the same residue numbering as in Hao *et al*., see supplementary file for explanation). Then a rotation of GA_4_ occurs to interact with TYR247 and ASP243 to allow chaperoning to more buried interactions with HIS119 and ARG244. Finally, GA_4_ resides in the binding pocket with its carboxyl group interacting strongly with SER116 and SER191. Excluding the two binding pocket serine interactions in the docked state, these six residues were identified by Hao *et al.* and are used here to identify the same pathway in our data using a large set of GA^(x)^.

For each GA variant, the set of GID1A residues that it interacts with when leaving through the channel are summarized in Table 2. In most systems the previously identified hydrogen bonding residues from Hao *et al.* are recurrent. It is notable that most of the hydrophobic contact residues are simply adjacent to these residues, but are highly conserved across the phylogeny of GID1. The only recurrent residue with large discrepancy between the GA^(a)^ and GA^(i)^ subsets is ASN32. Unlike the pocket residue interactions, the GA^(i)^ and GA^(a)^ interacted similarly with channel residues along the egressions. In simulations without the GAI N-terminus (gG systems), GID1A was found to be flexible enough to allow egression from this same pathway without a large portion of the interactions reported in Hao *et al.*, Table S1. Residue interactions we identified in the channel pathway in both the gG and the gDG systems were THR240, ASN32, and MET220.

**Table 2:**
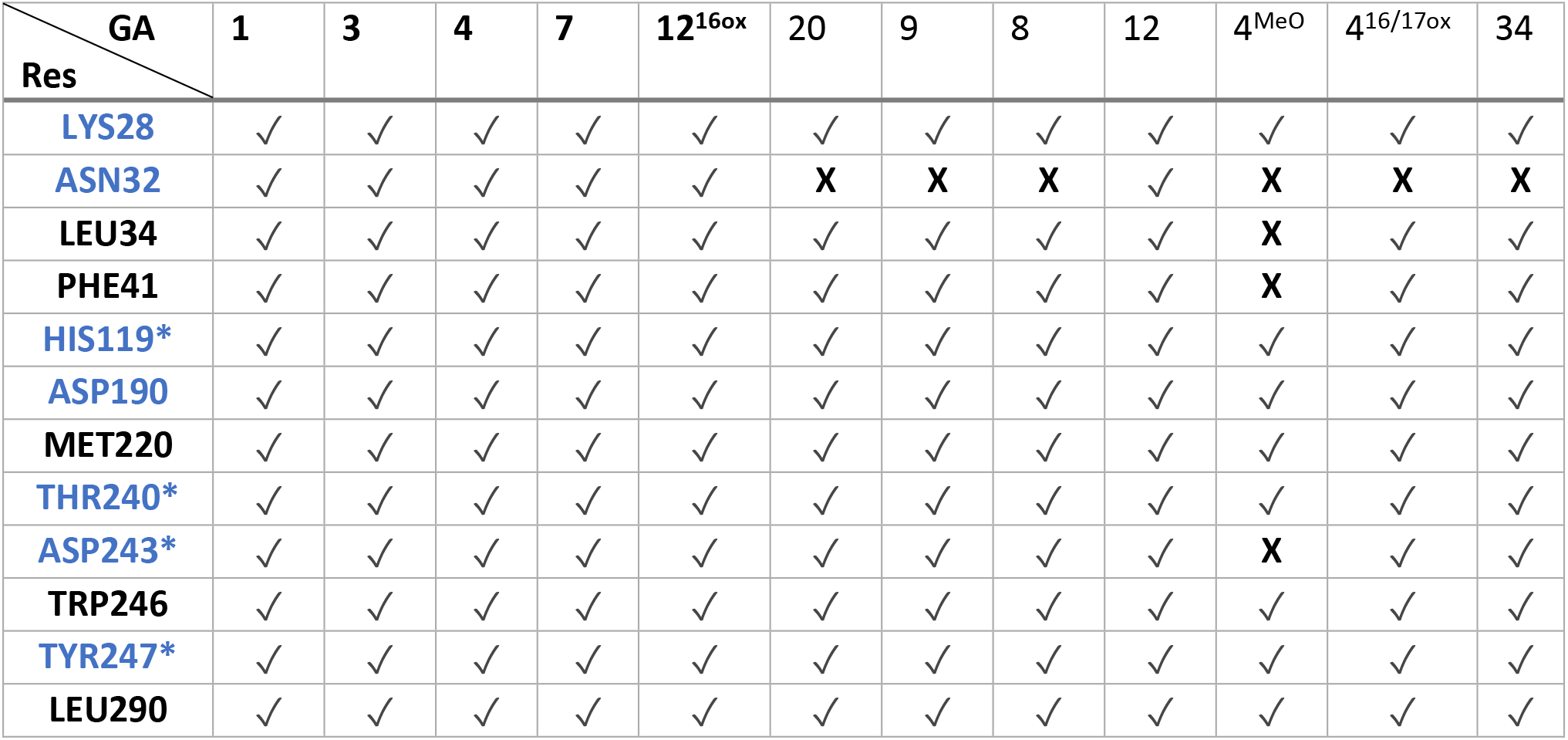
Hydrogen bonding and contact recurrent residues during Channel pathway egressions gDG (* denotes reported in Hao et al. as important pathway residues). Recurrent hydrogen bonds, in blue, and hydrophobic contact residues, in black, during production runs of the gDG system. GA variants in bold are reported bioactive. Recurrent set generated from GA^(a)^ subset and any pocket residue reported in Table 1 is removed for clarity, and otherwise, **X** denotes absence of interaction in given system, while check denote presence of interaction.

A second pathway is also uncovered, as shown in Figure 3 in blue. Without the DELLA protein, this pathway is slightly shorter than the channel even with dynamic conformations considered, *ca.* 6-8 Å vs. 9-12 Å. This cleft pathway is also more flexible than the channel with a cross-section of 8 Å that can expand up to 14 Å, versus the channel with 9 to 11 Å during egressions. Noting the differences in the blue egression plots between the top and bottom panels in Figure 3, the addition of the DELLA subunit causes a major change to the cleft egression path. While the addition does not heavily perturb the channel pathway, the cleft pathway must egress closer to the N-terminal of GID1A near MoRF 1 of GAI. The cleft pathway is visualized in Figure 4 and Figure S2. The channel pathway requires rotation around the shorter side of the GA molecule, while this cleft requires only minor rotation to align the molecule. Assistance with this alignment is offered from the electropositive interior of the pocket by attracting the carboxylate moiety and position the GA to bind into the pocket. Both the matching shape and electrostatics show a clear binding pathway, which is a persistent signature in all the simulations. Movie S1 and S2 provides a dynamic comparison of the egression pathways and their essential dynamics through the channel and cleft pathways, with and without the DELLA protein.

**Figure 4:**
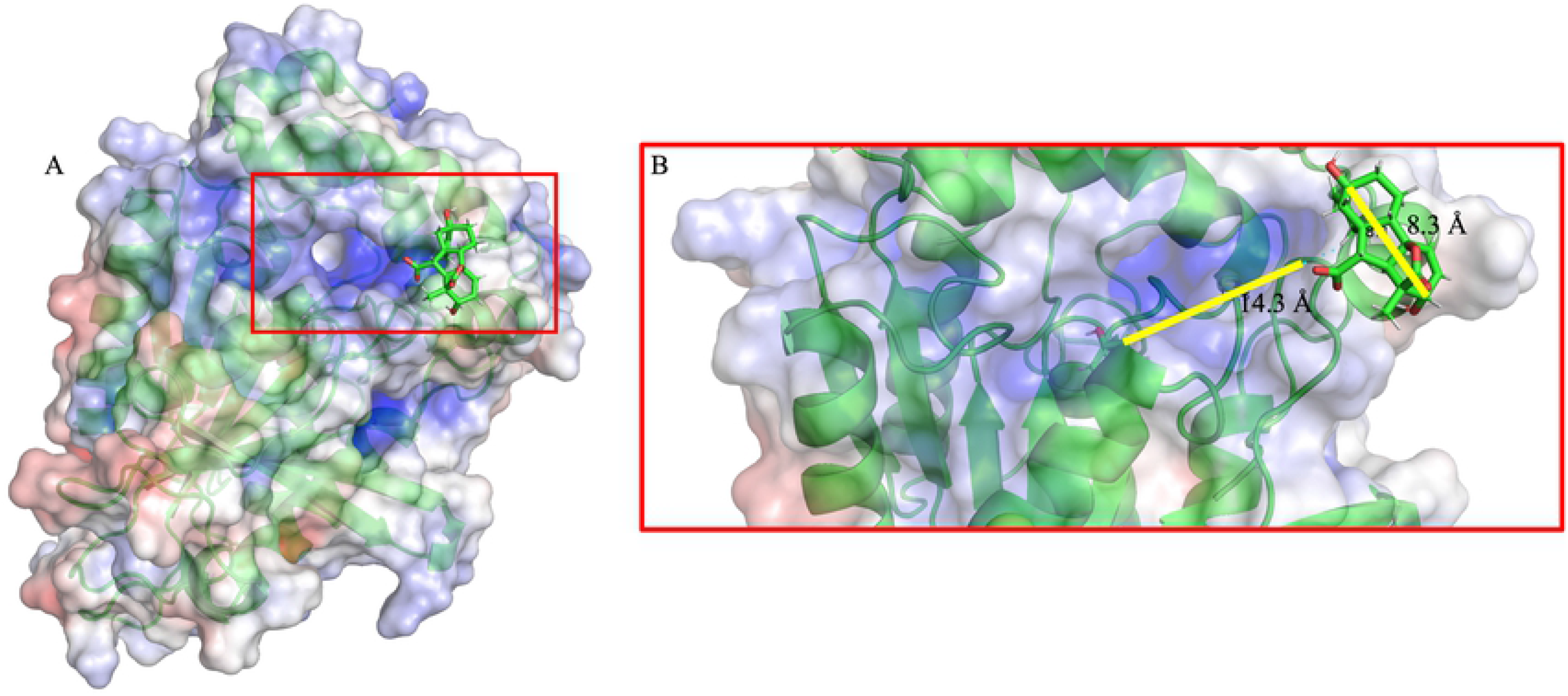
(A) Electrostatic surface, Pymol APBS plugin, of GID1A in the last frame from an egression via N-terminal cleft pathway. (B) Comparison measurements of a GA molecule compared to the cleft opening.

When the DELLA protein is bound to GID1A, the cleft opening is blocked. Nevertheless, as Figure 3 indicates, GA^(x)^ can egress from the cleft pocket opening even in the presence of the DELLA protein, albeit in a heavily perturbed pathway. The binding pocket residence times for the set of GA variants considered here for the gDG system are summarized in Table 3. On average the egression pathways through the channel is of higher propensity than the cleft pathway. However, among the 12 GA variants considered here, GA_9_, GA_4_ and GA_34_ are exceptions that exhibit a higher propensity to leave the binding pocket through the cleft pathway. The dipole moment calculated for each molecule displays little correlation to average residence time, as many of the GA^(a)^ share very similar values.

**Table 3:**
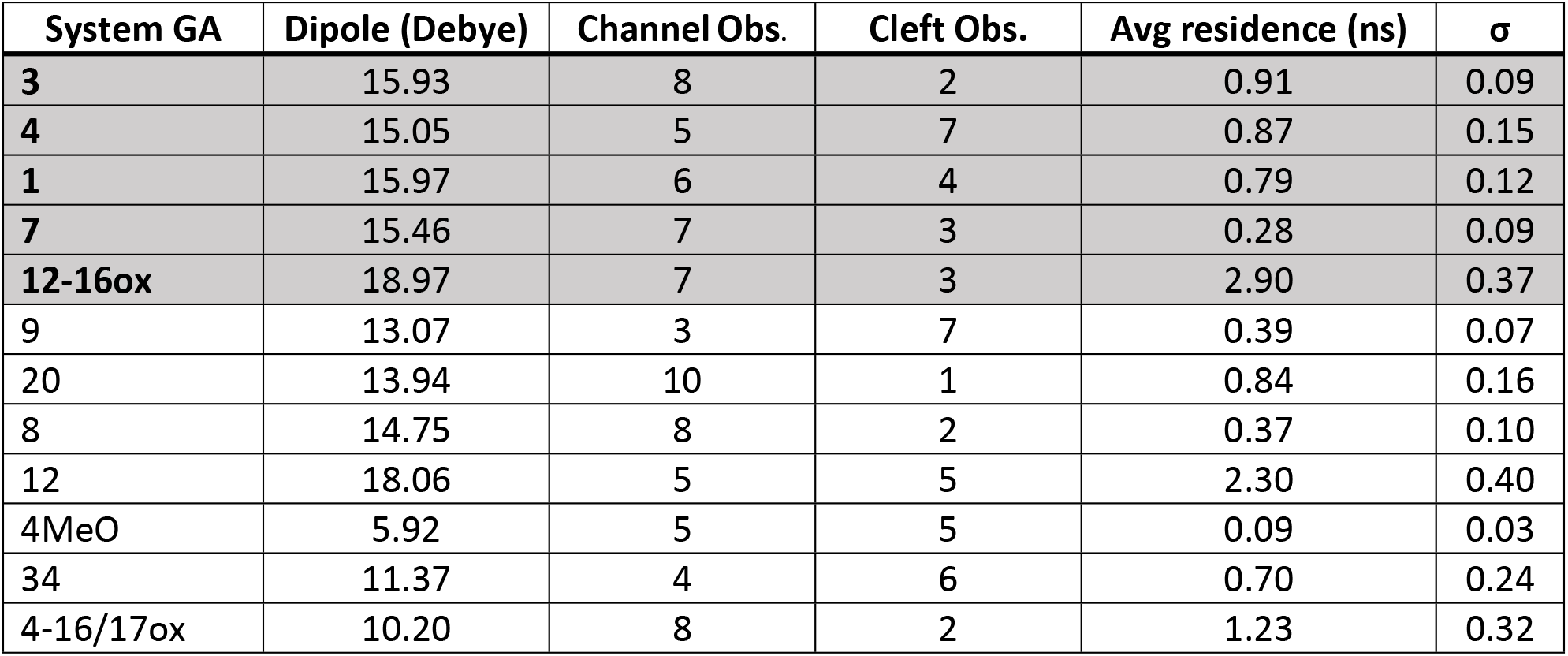
Residence times of GA in gDG and observed pathways from tRAMD egression experiments. Known GA^(a)^ data are shaded grey. Average residence time of total egressions for given GA^(x)^ gDG system, with the standard deviation also reported.

| System GA | Dipole (Debye) | Channel Obs. | Cleft Obs. | Avg residence (ns) | $\sigma$ |
| --- | --- | --- | --- | --- | --- |
| **3** | 15.93 | 8 | 2 | 0.91 | 0.09 |
| **4** | 15.05 | 5 | 7 | 0.87 | 0.15 |
| **1** | 15.97 | 6 | 4 | 0.79 | 0.12 |
| **7** | 15.46 | 7 | 3 | 0.28 | 0.09 |
| **12-16ox** | 18.97 | 7 | 3 | 2.90 | 0.37 |
| 9 | 13.07 | 3 | 7 | 0.39 | 0.07 |
| 20 | 13.94 | 10 | 1 | 0.84 | 0.16 |
| 8 | 14.75 | 8 | 2 | 0.37 | 0.10 |
| 12 | 18.06 | 5 | 5 | 2.30 | 0.40 |
| 4MeO | 5.92 | 5 | 5 | 0.09 | 0.03 |
| 34 | 11.37 | 4 | 6 | 0.70 | 0.24 |
| 4-16/17ox | 10.20 | 8 | 2 | 1.23 | 0.32 |

Given that the GAI MoRF 1 blocks the cleft, the tRAMD method was applied to the gG system for a subset of GA^(x)^ when the cleft is open. It was found that the GA^(x)^ have a choice of egressing out of the binding pocket through the cleft or the channel. The residence times found for the subset of GA^(x)^ with respect to the gG system are summarized in Table 4. Across the GA variants considered, on average, there is greater propensity to egress out of the cleft. However, GA_20_ and GA_12-16ox_ represent two exceptions. Therefore, it is clear that both egression pathways are viable in both the gDG and gG complexes for all GA^(x)^, with a variable degree of propensity for the cleft over the channel.

**Table 4:** Residence times of GA in gG and observed pathways from tRAMD egression experiments. Known GA^(a)^ data are shaded grey. Average residence time of total egressions for given GA^(x)^ gG system, with the standard deviation also reported.

| System | Dipole (Debye) | Channel Obs. | Cleft Obs. | Avg residence (ns) | $\sigma$ |
| --- | --- | --- | --- | --- | --- |
| **3** | 15.93 | 2 | 8 | 0.28 | 0.04 |
| **4** | 15.05 | 3 | 7 | 0.25 | 0.05 |
| **7** | 15.46 | 3 | 7 | 0.08 | 0.02 |
| **12-16ox** | 18.97 | 5 | 5 | 1.03 | 0.21 |
| 20 | 13.94 | 6 | 4 | 0.18 | 0.05 |
| 12 | 18.06 | 3 | 7 | 0.47 | 0.10 |
| 4MeO | 5.92 | 2 | 8 | 0.04 | 0.005 |

These data show that residence time alone does not offer a simple explanation for bioactivity. Even within the known bioactive set of GA ligands there is variation in average residence times particularly between GA_7_ and GA_12-16ox_. To glean a mechanistic explanation, each egression pathway was analysed separately where the counting of hydrogen bonds and hydrophobic contacts during production runs found that GA_7_ had significantly more hydrogen bonding to residues TYR247, GLY115, and VAL319. During these progression runs the GA_7_ ligand exhibited greater mobility than any of the other putative active GA^(x)^ ligands, in both the gG and gDG systems. A simple mechanistic explanation for markedly short egression times in GA_7_ could be differences in mobility within the pocket of the DELLA · GID1A complex due to the lack of an alcohol group at C13. However, no correlation was found between RMSD values and egression times across the GA^(x)^ to support such an explanation, Figure S5.

Analysis of the contact residues in the cleft pathway are shown in Table 5. In the complexed gDG system the cleft pathway is more complicated as the GAI MoRF 1 largely covers the cleft, but the space for this pathway is intrinsically present in the binding pocket of GID1A. As a GA^(x)^ leaves GID1A in the gDG through the cleft, it must directly pass the DELLA residues in MoRF 1 near GAI-LEU32, as highlighted in Table S2, and the nearby GID1A residue LEU217.

**Table 5:** Hydrogen bonding and contact recurrent residues during Cleft pathway egressions gG. Recurrent hydrogen bonds, in blue, and hydrophobic contact residues, in black, during production runs of the gDG system. Recurrent set generated from GA^(a)^ subset and any pocket residue reported in Table 1 is removed for clarity, and otherwise, **X** denotes absence of interaction in given system, while checks denote presence of interaction.

| GA<br>Res | 3 | 4 | 7 | 12<sup>16ox</sup> | 20 | 12 | 4<sup>MeO</sup> |
| --- | --- | --- | --- | --- | --- | --- | --- |
| LEU23 | ✓ | ✓ | ✓ | ✓ | ✓ | ✓ | ✓ |
| LYS28 | ✓ | ✓ | ✓ | ✓ | ✓ | ✓ | **X** |
| LEU49 | ✓ | ✓ | ✓ | ✓ | **X** | ✓ | ✓ |
| ARG51 | ✓ | ✓ | ✓ | ✓ | **X** | ✓ | ✓ |
| HIS119 | ✓ | ✓ | ✓ | ✓ | **X** | ✓ | ✓ |
| SER121 | ✓ | ✓ | ✓ | ✓ | ✓ | ✓ | ✓ |
| MET220 | ✓ | ✓ | ✓ | ✓ | **X** | ✓ | ✓ |
| LEU290 | ✓ | ✓ | ✓ | ✓ | **X** | ✓ | ✓ |
| ILE291 | ✓ | ✓ | ✓ | ✓ | **X** | ✓ | ✓ |
| GLY320 | ✓ | ✓ | ✓ | ✓ | ✓ | ✓ | ✓ |

### DELLA dynamics and tRAMD uncapping

As a complementary analysis to removing the GA ligand from the complex pocket, tRAMD egressions on the GAI N-terminus were also performed to explore the coordination of DELLA onto GID1A. This was performed for all gDG systems previously discussed including the apo DELLA · GID1A system (aDG). Important residue subsets described as MoRFs are known to be conserved features of the GAI N-terminus and considered to be responsible for recognizing and binding to GID1A [22]. The residues for MoRF 1 and 2 were included in the Hao modelling but the residues linking MoRFs 1 and 2 and beyond were missing from their model. Nevertheless, Hao *et al.* concluded there are differences between holo, with GA_4_, and the apo DELLA · GID1A complex at the incomplete protein interface. DELLA protein residues, most of which are within or near the later part of MoRF 1, were found in that study to change interaction by conformation changes induced when GA_4_ was present in GID1A. These residues interacted with GID1A N-terminal residues LEU9 and ASN32 on the alpha helix lid of GID1A. Although it was concluded that GA_4_ recognition leads to a conformationally stabilizing response, the previous work was mainly informed by the static protein structure from X-ray crystallography. In light of these critical differences we analysed both residence time (Table 6) and key contact residues for the MoRFs in tRAMD uncapping modelling experiments.

**Table 6:** Residence times of GAI N-terminus uncapping from various GID1A systems, with the standard deviation reported. Known GA^(a)^ data are shaded grey.

| System GA | Dipole (Debye) | MD eq. (ns) | Avg residence (ns) | $\sigma$ |
| --- | --- | --- | --- | --- |
| APO | 15.93 | 20 | 0.75 | 0.15 |
| **3** | 15.93 | 60 | 0.92 | 0.21 |
| **4** | 15.05 | 20 | 1.90 | 0.53 |
| **1** | 15.97 | 20 | 2.56 | 0.38 |
| **7** | 15.46 | 20 | 2.34 | 0.79 |
| **12-16ox** | 18.97 | 20 | 2.88 | 0.48 |
| 9 | 13.07 | 20 | 1.42 | 0.44 |
| 20 | 13.94 | 20 | 0.75 | 0.16 |
| 8 | 14.75 | 20 | 0.58 | 0.11 |
| 12 | 18.06 | 20 | 2.84 | 0.63 |
| 4MeO | 5.92 | 20 | 3.60 | 0.72 |
| 34 | 11.37 | 20 | 0.64 | 0.15 |
| 4-16/17ox | 10.20 | 20 | 1.01 | 0.29 |

The hydrogen bond and hydrophobic contact analysis was performed for the DELLA egressions as done for GA ligands. Cursory viewing of the full DELLA · GID1A complex shows MoRF 1 and 2 are largely responsible for stable binding to GID1A. MoRF 3 is above the entire complex, but note that this model is lacking a large flexible linker region between the N–terminus and GRAS domain, as well as the remaining GRAS domain. We acknowledge that it is possible that MoRF 3 could interact differently than observed in our simulations when the entire GRAS domain is modelled. With this caveat, we analyse MoRF 3 similarly to the other two MoRFs. The analysis of all three MoRF contacts during uncapping shows that the DELLA subunit has long-lasting hydrogen bonds to GID1 in gDG. As an exception, only MoRF 3 maintained its local secondary structure, and it loses intra chain interactions most rapidly. Since there is a long flexible linker between the GRAS domain and MoRF 3, the weaker interaction with MoRF 3 is expected.

The residues previously identified as important in GID1A LEU9 and ASN32 were found to be entirely conserved when analysing the hydrogen bond and contact interactions with MoRF 1 during production runs of the complex containing the GA^(a)^ ligands 1, 3, 4, 7, and 12-16ox. Hydrogen bonding to GID1A residues 9 and 32 were not statistically different between GA(a) · DG and the apo aDG systems. Comparing the interaction behaviour of the residues of MoRF 1 between GA^(a)^ and GA^(i)^, only LEU9 was found to have conserved contacts.

Analysing the uncapping of DELLA (Table 6), shows a weak correlation with the bioactive GA systems taking longer to remove the DELLA subunit. An average of 2.12 ns for the GA^(a)^ systems versus 1.55 ns for the GA^(i)^ variants, with APO having an average of 0.75. Notably GA_4MeO_ and GA_12_ significantly increase the average residence time of the DELLA protein for the GA^(i)^ category. During uncapping events the ligand pocket seemed relatively undisturbed, e.g. the ligand did not lose or gain many new interactions in all systems. For all GA ligands, the series of key pocket GA contact residues mentioned in Table 1 were recurrent in these uncapping simulations as well.

MoRF 1 also showed long lasting hydrogen bonds to ASN19, LYS28, ARG51, and ASN326 residues of GID1A. Interaction with the N-terminal of GID1A were found to be largely stabilized by hydrophobic interactions. While both of these interaction sets persist for a large number of the uncapping events, the interactions often break before the end of the DELLA egression. It is notable that the DELLA-GLU26 from the eponymous DELLA motif of MoRF 1 directly interacts with ASN19 of GID1A and was observed to be long-lasting during uncapping.

Analysis of MoRF 2 shows recurrent interactions of the active GA systems are largely intra chain interactions, suggesting MoRF 2 controls the tertiary structure of the DELLA subunit. MoRF 2 also maintains long-lasting hydrogen bonds to TYR48 and ARG51 of GID1A. Comparisons between the GA^(x)^ systems and the APO system hydrogen bonding of MoRF 2 showed inter chain residue interactions to GID1A residue ARG51 to interact significantly more in the APO system. GID1A’s ARG51 likely swings between MoRF 1 and 2 during normal dynamics.

MoRF 3 was expected to only interact with intra-chain residues within DELLA due to its position above the entire complex (Figures 1 and 5). Interactions of the main bioactive GA systems showed the expected intra-chain interactions to consist of highly localized residues, within plus or minus five residues of MoRF 3. However, between the gDG and aDG systems the APO system exhibited unique interactions with, MET13, ALA41, and ASP42 of the GAI N-terminus. These are significantly farther away than any interaction in the bioactive systems, suggesting MoRF 3 has a different conformation closer to MoRF 1 of DELLA in the absence of a GA ligand, as visualized in Figure 5.

**Figure 5:**
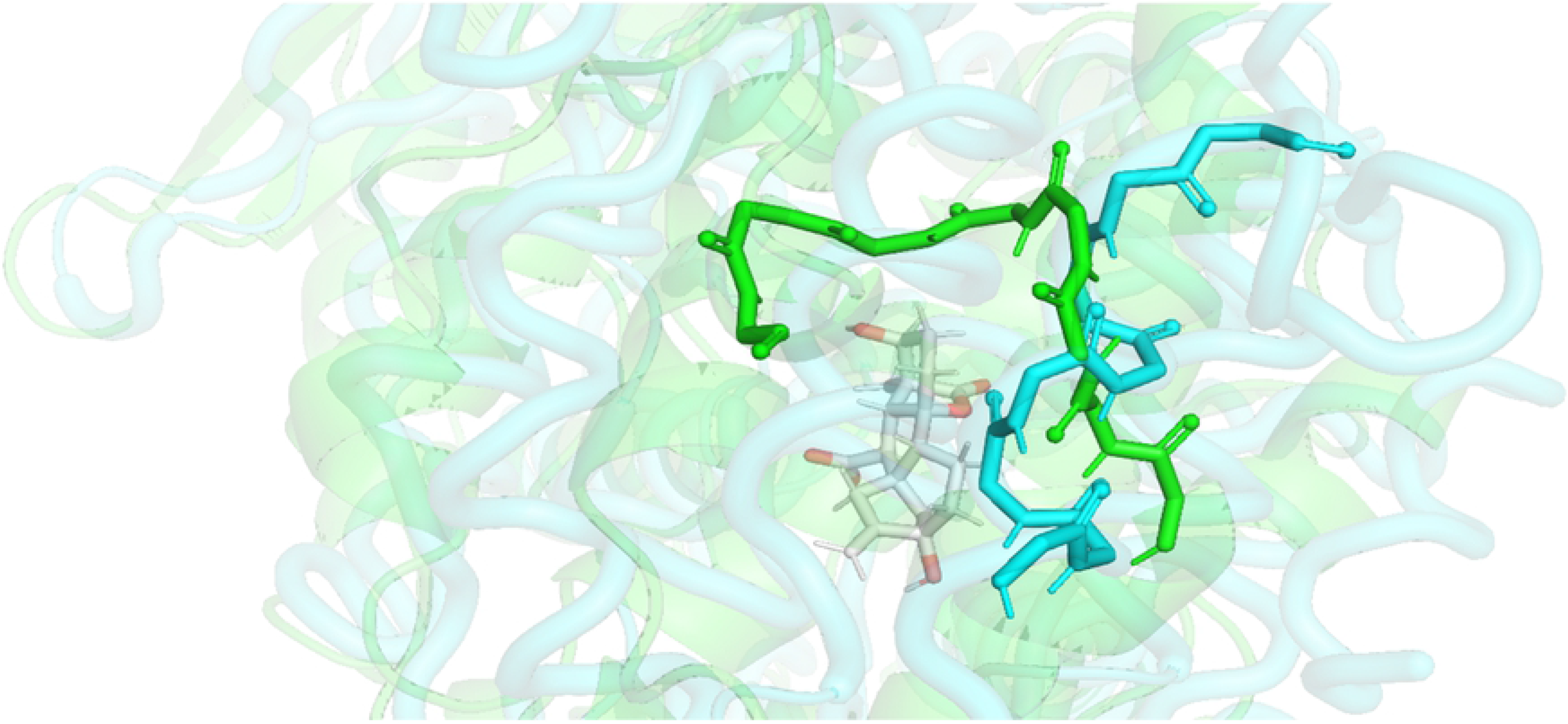
MoRF 3 conformations of typical active gDG, cyan, system compared with apo (aDG), green.

### Phylogeny of GID1

To examine the phylogeny of GID1 across a broad cross-section of plants from divergent taxa, we mined the existing database EnsemblPlants (34). Upon examining the phylogenetic tree for this family, it is clearly split into four major subtrees. These four clades consist of 1) basal Angiosperms and Moss ancestral plant species, 2) monocot GID1 (one major branch only with duplications in later parts of the phylogeny), and two major dicot branches 3) the GID1A and 4) GID1B branches – the latter two housing the Arabidopsis GID1A and GID1B members respectively. Comparing these trees with the plant evolutionary tree of life suggests several duplication events have occurred in the GID1A and GID1B branches (identified as duplication or ambiguous nodes in the tree). These factors were combined to derive 26 major subgroupings from which consensus sequences were created by setting conservation at 90%. This cut-off of 10% protein variation for proteins spanning the 50–100 million years in evolution of these suborders and superclasses is equivalent to 5–50 times lower per million years (/m.y.) when compared to the degree of conservation between alleles (0.1–0.2% /m.y. versus 2–3% allelic variation separated by short time periods of the order of a few million years at most and equating to approximately 1–5 % /m.y.). We suggest this cut-off is a conservative measure and identifier of key conserved residues across broad taxonomic groups of plant families.

We examined the conservation at 37 key GA^(x)^ contact residues identified in the computational analysis across the four main GID1 groups (basal GID1, monocot GID1, dicot GID1A and dicot GID1B, results detailed in Supplementary Tables 3–5). In summary 11 out of the 15 pocket contact residues show a high degree of conservation and the remaining 4 only show conservative substitutions, 10 out of 12 channel contact residues showed a high degree of conservation (some with a degree of conservative substitutions) and only 2 showed non-conservative substitutions while 8 out of 10 cleft residues show a high degree of conservation and the remaining 2 only show conservative substitutions. These results suggest a high degree of conservation across all of the GID1 residues that contact the GA ligand. The key GID1 cap residues (alpha helix A, B and C) in contact with the MoRFs of DELLAs are also highly conserved or have conservative substitutions (data not shown).

## Discussion

### The influence of cleft, channel and MoRFs on GA binding

The description for how DELLA GAI binds to GID1A has common aspects across all our models. First, the GID1A lid must be closed for DELLA GAI to bind to GID1A. In absence of direct experimental evidence, it is presumed that the DELLA GAI protein only binds to GID1A in the presence of GA. Once formed, the gDG complex subsequently attracts an E3 ubiquitin ligase (SLEEPY1 in Arabidopsis), a component of the ubiquitin-mediated machinery that initiates a programmed degradation of the attached DELLA protein. All prior work neglected to consider the effect of GA variants by assuming that only bioactive GA plays a role in the earliest part of this process. Here, we also considered the possible effects of inactive GA on the DELLA binding process, which is important for developing a complete kinetic model.

We found that the same application of force in tRAMD simulations produce egressions of GA^(x)^ from the gG and gDG systems. Not-withstanding exceptions, a general trend indicates there is higher propensity for cleft pathway egressions in gG and channel pathway egressions in gDG. Furthermore, any GA ligand present in the DELLA · GID1 complex is able to stabilize the GAI subunit in its attachment to GID1A. This is suggested by results from comparing uncapping residence times, finding that they are, on average, higher for all tested gDG systems than for the aDG systems. This result applies to each GA^(x)^ case studied separately. Taking this result alone, we cannot discriminate any critical differences between GA^(x)^ that would classify them as bioactive or inactive. This is not to say there are no differences. Consistent with our production run results, it is likely that the different GA^(x)^ create differences in conformational dynamics within the gDG complexes (not studied here) that affect the degradation of the attached DELLA protein.

Relatively lower RMSD was found in the MoRFs compared to the linkers between the MoRFs in both gDG and aDG systems. Overall mobility in the DELLA protein is less in gDG compared to aDG for each GA^(x)^. As Figure 2 shows, the region between MoRF 2 and MoRF 3 has greater mobility with known bioactive GA compared to known inactive GA variants. The RMSF for the carbon alpha atoms in the key interacting pocket residues of GID1A are found to be less mobile in the aDG and gG systems compared the gDG systems. In contrast, the DELLA protein RMSF has greater mobility in aDG than found within the gDG systems, see Figure 2 and Figures S8 and S9 for additional RMSF data. These comparisons were made across our GA^(x)^ set. Shifts in mobility from one region to another is easily understood by an enthalpy/entropy compensation mechanism driven by rigidity changes (2). Conformational entropy increases in certain regions that become more flexible, while other regions become more rigid with more favourable enthalpy due to conformational shifts in a protein. The greater RMSF found in the DELLA protein when complexed in the aDG system is consistent with experimental data that without GA the DELLA protein does not bind to GID1A. In essence, any of the GA^(x)^ forms can act as a cofactor. Furthermore, if GA leaves gDG, it is expected that aDG becomes an unstable structure with a relatively short lifetime relative to the gDG lifetime. However, our data show it has a relatively long lifetime relative to GA binding to aDG through the channel pathway. These examples show that the DELLA protein modifies the inherent dynamics of GID1, with details that depend on the GA variant bound to GID1 and whether it is in apo form.

### Sequence of events and bioactivity of Gibberellin

Determining the bioactivity for GA variants is an ongoing challenge of exploration, as illustrated by the recent addition of DHGA12 into the bioactive category (33). With this line of classification, there is an underlying assumption that appropriate concentrations of specific GA variants must be present at certain plant tissues or cells. An intricate web of GA modifying enzymes exists and these are clearly differentially regulated under particular environmental stimuli (35). These modification systems presumably shift GA^(x)^ population abundance at particular cellular locations. However, a mixture of GA variants will likely be present at most locations. Unless there is an overwhelmingly strong differentiation between bioactive and inactive GA, the binding process that takes place to form gDG should account for competitive binding. Our results show that residence times of GA variants do not correlate to bioactivity. There is a trend in our data that residence time increases when the GA moieties support more intra-molecular H-bonding. These results suggest there are subtle mechanisms in play that govern the observed specificity in plant growth regulation. To understand these differences requires looking at a specific GA moiety along with propensities for the GA variant to form H-bonding and hydrophobic contacts within the cleft and channel pathways, which will affect association and dissociation rates.

In the absence of GAI, entry for any GA^(x)^ via the cleft pathway is observably more accessible than via the channel pathway. This is demonstrated by the fewer hydrogen bond interactions in cleft egressions in Table 5 when compared to channel egressions in Table S1 in the GID1A system. Along with the easier arrangement of GA carboxylate to organize into the pocket and the higher likelihood of observation of cleft egression (see Table 4), this suggests an easier route in the absence of DELLA motif. However, when GAI is bound to GID1A, by MoRF 1 and 2, the contact residues to GA at the surface of the cleft pathway and the side chain flexibility needed to open the cleft pathway are hindered. Egressions along the cleft pathway were still observed with GAI present but drastically altered. In the complexed state GA was forced to take paths directly under the DELLA MoRF 1 segment (see GAI-LEU32 contacts in Table S2). This suggests that GA^(x)^ egressions from the closed cleft will destabilize the gDG structure, as hydrophobic interactions at this interface are disturbed. This in turn could help control the necessary GA^(a)^ recognition, as the more bioactive GA ligands are less likely to leave or to interact with this aliphatic region. MoRF 2 interactions are found to be similar between the GA^(a)^ and GA^(i)^ systems, removing it as a possible source for discrimination. We also note that a number of the channel contacts evident in GA^(x)^ egression pathways are missing when the ligand is GA^(i)^. Thus, there is a higher propensity for GA^(i)^ to be expunged from the closed cleft than a GA^(a)^. Dynamical differences in the intra-chain interactions within DELLA are observed when bioactive gDG and aDG are compared.

The opening of the cleft pathway in GID1 is a hole that can fit a GA^(x)^ shaped molecule, and it forms when the unstructured loop that LEU323 is attached to becomes part of a partial helix. This releases helix B of GID1 to tilt outward as it lacks interaction with that loop (see Fig S2). This motion appears as a natural breathing motion within GID1 when the DELLA protein is absent. It is worth mentioning that based on structural similarities, this cleft might be the remnant of the clade IV carboxylesterases (CXE) binding pocket in ancient plants (36). Assisting interactions with HIS119 allow the GA^(a)^ ligands to twist, having the alkyl backbone face LEU323 and the aliphatic residues adjacent. These events were observed as essential dynamics of this localized region based on a principal component analysis. In Figures S3, S4, S6, and S7 we compare the influence of DELLA binding on these essential motions (37). From analysing the pocket residues that contact the ligand when fully docked, it is likely that key interaction to TYR322 prevent the LEU323 loop from opening up and allowing a facile egression. This tyrosine may also be important for bioactivity because it directly interacts with the key hydroxyl moiety on C3 of GA.

Based on the observation that the cleft opening is relatively large with a shorter passage to the binding pocket compared to the narrower and longer channel passage, we suggest GA^(x)^ binding to GID1A through the cleft in absence of DELLA protein is a significantly more rapid process than through the channel. The GAI is assumed to bind to gG with little distinction in GA^(x)^. Although it has been shown that mutations in GID1B can be made to facilitate GAI binding without the presence of any GA, this has not been observed in nature, suggesting the role of GA is to act as a cofactor to facilitate the gDG complex to form. In other words, it is not the case that an aDG complex cannot be stable, but rather, it is important for biological function to have the presence of GA as a facilitating condition in order to control growth regulation in a plant. Furthermore, GA^(x)^ dissociates from gDG on a much faster time scale than GAI can dissociate from GID1, implying there will be some degree of binding of DELLA to GID1 in the absence of GA.

### Proposed rectification mechanism that can select for bioactivity

We propose a novel kinetic model based on the newly found cleft pathway that also takes into account the role played by the DELLA N–terminus that includes three MoRFs. Our kinetic model elucidates how this system recognizes bioactive GA variants under competitive binding. Furthermore, we performed a phylogenetic analysis that shows there is conservation of key residues within the binding pocket, cleft and channel pathways – suggesting that the same general mechanism in the GAI-GID1A interaction will apply to all DELLA · GID1 complexes.

There is uncertainty in whether a particular GA is bioactive for a given receptor. Therefore, we introduce a more appropriate language to describe GA variants in terms of major and less active forms of GA^(x)^ relative to a given receptor. Subject to a wide spectrum of activities, GA^(a)^ denotes a subset of major active forms, while GA^(i)^ denotes a subset of less active forms. For simplicity of this discussion, the two subsets represent extremes in the activity spectrum for the GID1A receptor (active versus inactive). The rank order of GA variants from most bioactive to inactive would likely be different with respect to a different receptor. If the sequence of another receptor is highly conserved with respect to GID1A in the binding pocket as well as the two binding pathways, then the rank ordering of GA activity will presumably be similar. To emphasise generality of the kinetic model, we discuss its components in terms of the GAI and GID1 family of proteins.

Our results suggest that a molecule resembling the columbic and geometrical properties of GA^(x)^ can bind into GID1, particularly via the cleft pathway. Notably this includes GA variants that are often only described as “inactive” or “intermediates” in the literature. Therefore, a degree of caution should be exercised when classifying GA^(x)^ as bioactive or not. With this in mind it is axiomatic that recognition of bioactivity must involve GID1 in some capacity. If GID1 can bind an array of GA^(x)^ that likely includes both bioactive and inactive examples, then, this suggests a mechanism of action removes inactive (or less active) forms of GA from the receptor or affect the DELLA protein binding in other ways. Differentiation between GA^(a)^ and GA^(i)^ can occur before and/or after DELLA is bound to the gG system. As shown in Figure 6, we construct a generic 1st order kinetic model for a proposed rectification process.

**Figure 6:**
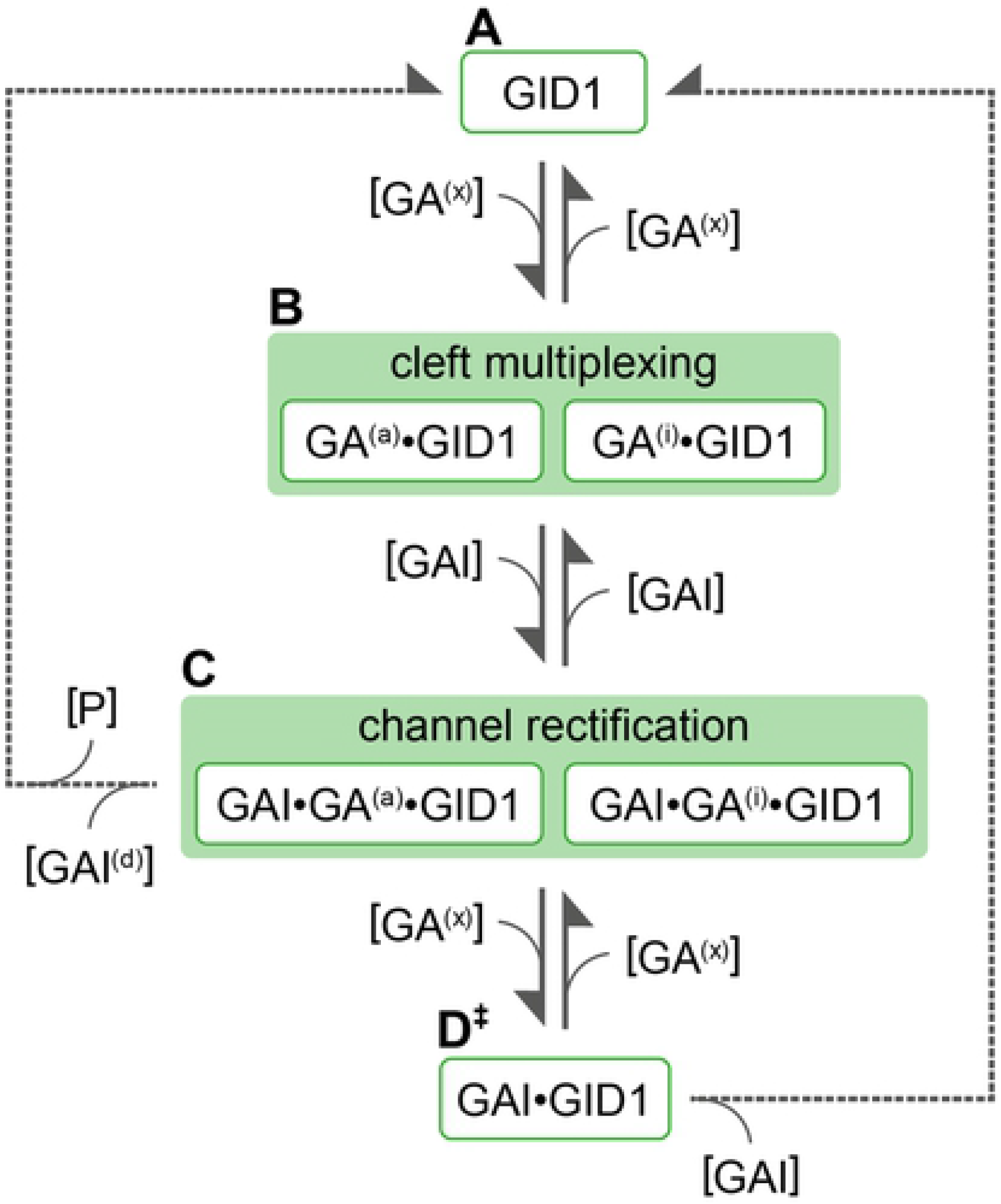
Synergetic three-stage kinetic model of DELLA protein assisted rectification of GA in GID1A receptor.

The GA^(x)^ in Figure 6 refers to any GA variant, but once a GA variant interacts with GID1, it then becomes possible to classify the GA as a major or less active form. As a simplification, only the extremes are tracked to show the possibility for competition between active and inactive forms of GA, denoted as GA^(a)^ and GA^(i)^ respectively. This simplification demonstrates salient features of the model by considering kinetic rates for only two types of GA (active and inactive), but in reality, different kinetic rates will likely apply to each GA variant comprising a broad spectrum of activities.

In state A, we have a free GID1 in an environment containing a mixture of GA variants, where GID1 has opportunity to bind to any GA variant, represented by GA^(x)^. Since the open cleft does not have a high level of specificity for the type of GA^(x)^ to bind, the cumulative concentration of all GA variants drives GA association to state B. The GA · GID1 complex is now primed to bind with GAI. Since GAI association is a relatively slow process, it is expected that many cycles of association and dissociation of different GA variants through the open cleft in GID1 occurs in the interim. The key role of the cleft opening is to multiplex the kinetics of all GA^(x)^ associations that can take place to populate state B while depleting state A as much as possible depending on a variety of details. While details depend on environmental conditions such as concentration of repressor, receptor, protease, and GA variants, the qualitative features remain robust. Eventually, GAI binds to GID1 to form the GAI · GA · GID1 complex that defines state C.

The details of which GA variant is bound affects the stability and conformational dynamics of the GAI · GA · GID1 complex, which in turn affects the propensity for the E3 ubiquitin ligase to exercise proteolysis to degrade GAI. The GAI · GA^(a)^ · GID1 complex is primed for GAI to undergo degradation, but the GAI · GA(i) · GID1 complex with less active GA varaints is unsuitable, being the bottleneck that ultimately differentiates the bioactivity across GA variants.

Dissociation of GA variants is possible through the closed cleft (the cleft is blocked by GAI) and the open channel within GID1. Since the time scale of GAI dissociation is relatively long compared to GA dissociation and association from the binding pocket in GID1, it is possible to reach state D.

State D is not stable. Eventually the unstable GAI · GID1 complex will dissociate because a GA variant is necessary to maintain its stability. However, this dissociation process takes place on a time scale that is much longer than the time scale for GA variant association, mainly through the channel. The rates of dissociation and association will be nuanced depending on the moieties of the GA variant and the sequence of GID1, and especially the structure and dynamics of the GAI GID1 complex that also depends on the sequence of GAI (or indeed other DELLA proteins where they exist). The upshot of these details is that the overall dissociation rates for GA^(i)^ are greater than GA^(a)^ and the overall association rates for GA^(a)^ are greater than GA^(i)^ in the kinetics that bridge states C and D.

A kinetic equilibrium is set up that drives the concentration ratio of Σ*_a_*[GA(a)] to Σ*_x_*[GA(x)] toward 1. This result is insensitive to model details (i.e. the rates) provided the general trends described above are satisfied. As a consequence, rectification of less active GA variants takes place to amplify the propensity of the biologically important GAI · GA^(a)^ · GID1 complex. That is, the GAI · GA^(a)^ · GID1 complex will be well populated even within an environment where the ratio of Σ*_a_*[GA(a)] to Σ*_x_*[GA(x)] for the mixture of GA variants in solution is small. An amplification factor, defined by the ratio of Σ*_a_*[GA(a)] to Σ*_x_*[GA(x)] within the binding pocket divided by the ratio of Σ_a_[GA(a)] to Σ*_x_*[GA(x)] for the GA variant mixture can easily reach up to 10.

It is also possible that the dissociation rate for GAI from the GAI · GA^(i)^ · GID1 complex is greater than the dissociation rate for GAI from the GAI · GA^(a)^ · GID1 complex. This is another nuanced way rectification can take place. We note that this would be a significantly slower process than GA^(i)^ leaving the complex. Our data suggest both types of nuanced mechanisms can work in tandem. If both mechanisms are present to rectify the less active GA in the binding pocket, amplification factors near 100 can be obtained. Since we cannot offer any biophysical/chemical rationale for why these proposed mechanisms would not be present, we propose that it is through protein evolution that repressors and receptors become in tune with one another through a specific set of GA variants. Given the simplicity in the explanation of how a repressor-receptor pair functions with high fidelity, while making use of a large diverse set of GA growth hormones, it is fitting to look at evolutionary consequences in plant biology.

### Implications for evolution of the Gibberellin signalling system

The high degree of conservation over the 37 GA contact residues across the four main GID1 groups suggest that most of the conclusions we drawn about the ligand migration pathway from our analysis of GID1A are likely to be true for most or all other GID1 proteins. Out of this set of residues there were only two residues in the channel pathway that show a significant degree of substitutions that are not necessarily conservative in nature. The proposed kinetic model is quite plastic, where we suggest that the control of major and minor active forms of GA is best understood by an evolutionary perspective.

One of the more variable sites among the contact residues is an ASN residue 32 in GID1A that undergoes H-bonding with alcohol moieties of all bioactive GA during channel egressions and that can change to other polar residues (HIS, LYS, THR) or even a non-charged MET in basal species. A more detailed examination of these different forms of GID1 would be required to ascertain if it is likely that these changes could significantly change the behaviour of one or more GA variant as it passes through the channel. This residue is part of alpha helix B which appears to play a significant role in the receptor “breathing” motion that we discuss above. We suggest these variations can have significant impact on different versions of GID1 e.g. they may have somewhat different specificities for different GA variants or interact differently with the DELLA protein that also interacts with this alpha helix. Interestingly this ASN residue appears not to be contacted by many of the GA^(i)^ variants for which there is no convincing data for bioactivity. This could suggest the potential for differences in this “breathing” motion in some GID1 loci associated with particular GA variants and perhaps even a lack of bioactivity in the context of a particular gDG complex.

The second contact residue showing a significant amount of non-conservative substitutions is a THR residue in GID1A-240 that was also identified by Hao *et al*. as a key GA contact residue and is a residue that apparently undergoes hydrogen bonding with GA lactone. This residue partners with ASP243 to initiate the channel pathway before GA^x^ rotates to interact with TYR247 and ASP243. The variability here seems to be large confined to the two dicot subgroups (GID1A and GID1B). This THR residue site displays substitutions to LYS, ARG and GLY which could well have major implications for H-bonding to one or more GA variant and being at the start of the pathway this could significantly impact on the rate of engagement of this pathway by different GA^(x)^ forms for GID1 variants carrying these substitutions.

The dicots are unusual in showing a very deep phylogenetic split into two major GID1 clades. Monocots generally contain just one GID1 locus or, if there has been any duplication, these are likely to be more recent events. The presence of two GID1 clades in dicots that have been evolving separately (but present together for a long time) would provide opportunities for specialized functions to evolve between these clades. Possible areas of partition of components of their combined function could include specialization for different GA variants, different degrees of GA signal transmission, or perhaps forming specialized interactions with a subgroup of DELLA proteins. It is notable that there are often multiple members in the DELLA family in dicot plant species as well.

In plant taxa where there is only one GID1 this specialization would not be possible and the residue of choice may be more of a compromise – perhaps explaining why this residue is highly conserved in the monocot and basal plant lineages and that there is a tendency for just one partnering DELLA locus to exist in monocots as well. Perhaps the ability of some GID1B proteins to interact with DELLA proteins in the absence of GA ligand (albeit it usually less strongly than in the presence of GA), similarly reflects some of this specialization of function. In addition to the major divide between GID1A and GID1B there have been a number of subsequent duplication events in both of these clades that predate the split between major divisions within the Asterids and Rosids. We suggest that a very early duplication event in the dicots released additional evolutionary potential for variation to occur without a loss of function. As a result, dicots may have taken advantage of this to evolve a more complex mix of GID1 and partnering DELLA loci. We would therefore expect there to be a higher degree of specialization in these loci not possible in monocots and basal plant species. The higher variability in two GA^(a)^ contact residues in the channel pathway of dicots may also reflect this higher propensity for specialization in dicots.

## Conclusion

Based on data from simulation, experiment, and evolutionary analyses, we put forth the most complete kinetic model for how gibberellin promotes plant growth by enhancing binding between GAI repressors and GID1 receptors to regulate GAI degradation and control plant growth by de-repressing GA response genes. There are two binding pathways in the form of a cleft and channel in GID1. The cleft pathway is open when GAI is not present, giving easy accessibility to GA variants to bind within the binding pocket. The GA variants acting as cofactors help GAI bind to GID1. After the formation of this complex, if the GA variant is bioactive, a subsequent process for the degradation of GAI eventually follows. However, there is a continual kinetic process that takes place with GA variants leaving and re-entering the binding pocket. Due to large differences in time scales, many cycles of GA exchanges can take place before the GAI · GID1 complex has time to dissociate, which is inevitable if no GA re-enters the binding pocket. As major and less active forms of GA enter and exit the binding pocket recurrently with disparate rates that favour the major active forms, rectification of less active forms occurs to yield higher propensity for a major active form to control growth.

The biological ramification of this kinetic model partly accounts for the highly diverse set of GA variants known to be present in plants. Within the cellular environment there will be a mixture of GA variants available to bind to GID1 through competitive binding. In the initial binding of GA to GID1, the consequence of binding of major active or less active forms of GA is similar. The binding of GAI to GID1 will be enhanced for almost all GA variant. The interactions between GAI and GID1 in the GA · GAI · GID1 complex further promotes specificity. If the captured GA is a major active form, the degradation of GAI can take place via the known SLY1 facilitated ubiquitination and proteolysis. Otherwise, the rectification process takes place, which is especially needed in cases where the concentration of the major active form is considerably less than the combined set of less active forms. This mechanism allows plants to readily sense fine concentration changes of bioactive GA molecules with high specificity to enable certain GA variants to trigger specific signalling events. In this scenario, the large number of different GA variants found in plants becomes biologically advantageous for homeostasis and/or specialisation when combined with different DELLA · GID1 complexes (particularly in dicots). Our proposed mechanism is consistent with the general paradigm of protein-ligand binding and protein evolution. Overall, passive specificity is enabled by small changes in pathways of GID1, which gives rise to the complex series of interactions with the disordered DELLAs in a variety of plants which enables their complex growing habits.

## Methods

### Computational modelling and analysis

The Murase *et al*. (27) resolved structure from *A. thaliana* GA receptor GID1A and the N-terminal fragment of the GAI conserved across various plant species (PDB Accession no: 2ZSH) was partially unresolved. This is explained by the inherent disordered nature of the GAI N-terminal structure (22). The absence of large parts of the disordered domains prevent realistic simulation of this hormone receptor system. Therefore, missing segments were constructed using I-TASSER (Iterative Threading ASSEmbly) (28) ab-initio and threading method software. The 110 residue N-terminal DELLA sequence from the 2ZSH structure was used to generate a more complete model by filling in the missing components between the three MoRF segments that were partially predicted in 2ZSH.

The DELLA structure represents a designed subsection of the DELLA GAI sequenced used in the source study. As such the source material sequence published with 2ZSH (chain B) is used to number these residues (1–110). The GID1A (chain A) structure in this model is missing 5 residues from the N-terminal and 2 from the C-terminal ends. Both 2ZSHA and 2ZSHB have a short remnant of an artificial HRV-3C protease cleaved His-tag at the N-terminus. For GID1A the residue numbering includes these missing residues as aligned from uniport *Arabidopsis thaliana* sequence for GID1A (UniProtKB: Q9MAA7-1). We exclude residues in the tag remnant from further discussion.

Ten full atomistic models of the DELLA disordered protein were generated from residues 1 to 110. Variation in the top models at the disorder loops showed minor orientation differences. The top scoring model was checked by potential energy calculations using ProSA-web (38). ProSA is based on PDB structures derived from X-ray diffraction (XRD) and Nuclear Magnetic Resonance (NMR) based PDB structures to score unseen structures. The deduced DELLA structure is consistent in conformation to NMR structures and consistent with Ramachandran backbone angles. To arrive at the gDG system, the DELLA GAI N-terminus was placed by a structural alignment to the original fragments of the DELLA subunit. Furthermore, each GA^(x)^ ligand was aligned to the 2ZSH structure before minimization for both gDG and GA^(x)^ · GID1A (gG) systems. All GA that had carboxylic C6 were deprotonated before calculating ligand parameters or docking into GID1A (39). All simulations were carried out on Linux based high performance computing clusters.

The GA variants were optimized in AMBER using semi-empirical BCC available inside the Ambertools software (40), then the ligands were aligned with GA_3_ of the crystal 2ZSH before minimizing the system in NAMD. In each system solvent within 5 nm of the 2ZSH GA_3_ was kept and placed back into the structure before energy minimization as defined in Kokh *et al.* (31). The steps performed in this process consisted of the preparation of the solvated protein-ligand complex, followed by a 20 ns production run, or a steady state simulation, of water and NaCl ions at 0.5 mmol with the various systems studied at 300°K, followed by multiple tRAMD simulations in NAMD to obtain ligand dissociation trajectories. The tRAMD relative residence times were obtained using a previously reported protocol (31) adapted to Matlab using PDFEstimator (41) in Matlab 2020a. Systematic evaluation of force constants found that at a force of 20 kcal mol^−1^ Å^−1^ allowed egression to a threshold of 30 nm from the docked position in a computationally tractable timespan. Similar force evaluation was performed for the egressions of the entire DELLA protein. A significantly higher force of 80 kcal mol^−1^ Å^−1^ was found to allow egression on ns time scale to reach a distance of 30 nm from the original position of DELLA chain.

Visual Molecular Dynamics (VMD) analysis of trajectories was performed using in-house designed scripts in TCL. A cut-off distance of 3.5 Å and angle of 30° for hydrogen (H) bond counting. Hydrophobic contacts were defined using known nonpolar residues and a contact cut-off distance of 3.5 Å for conservative estimates of contacts. These were tabulated into a text file for further processing. Raw counts of interactions to residues from a defined reference set were used as samples in statistical analysis. Significance of differences were selected from a rational p-value threshold of 0.05. Recurrent residue sets were simple logical comparisons between lists of interaction residues between various simulation sets. For recurrent residues in various important comparisons the active GA subsets were used to create a residue subset, which was then compared to inactive GA interaction lists.

### Phylogenetic analysis of Gibberellin signalling components

We subjected both the N-terminus of DELLA proteins (up to but not including the GRAS domain) and the GID1 receptor family to a similar phylogenetic analysis in order to determine if residues of interest from the above analyses were broadly conserved or variable. The degree of variation within the disordered segments of the DELLA family N-terminus in particular only re-enforced that the three MoRF regions identified previously by analysis of a smaller dataset (4) were highly conserved. Most of the residues identified in the above modelling analysis were present in these conserved MoRF motifs. As this phylogeny has little new significant data, it is not presented. In contrast the GID1 protein family is clearly much more highly conserved across its entire protein sequence. Gene trees developed under Plant Compare of EnsemblPlants were used as a basis for obtaining the orthologs of the GID1 family of proteins from all plant taxa and their basal ancestors (Basal Angiosperms like *Amborella* genus and Lesser club moss sequences from *Selaginella moellendorffii*). There are no sequences in the more basal Earth mosses or liverworts (*Physcomitrella* and *Marchantia* genera). This tree contains four major subclade branches.

The members of four main subclade branches (basal plants and mosses, monocots and two major dicot subclades) were further split into major divisions at the suborder (monocot) and superclass (dicot) levels, combined with evidence for inferred duplication events in the GID1A and GID1B clades. Based on this analysis the dataset was divided into 26 groupings with 1 to 22 member proteins. The members of each grouping were then downloaded as a set of aligned groupings into Geneious, given a new nomenclature to identify the groupings and realigned in Geneious with default settings.

These alignments were examined for proteins with large gaps in the core of the GID1 sequence and N- or C-terminal extensions. Proteins with large deletions in the core were completely deleted from the subsets as these are likely to be miss-annotated and this will affect the conservation level of residues in the deleted regions. Large unique N- and C-terminal extensions were removed from the alignments as they result in extensions with apparent “perfectly conserved” protein regions due to the lack of other aligned sequences – thus leading to misleading conservation information. A consensus sequence set at 90% conservation was derived from each of the modified 26 groupings, and named via the new nomenclature. These consensus sequences were then re-aligned as subtree alignments and also the full set of all plant consensus proteins.

The aligned sequences are ordered with the most basal consensus group at the bottom and more derived consensus groups (as inferred from the EnsemblPlants tree) progressively aligned towards the top of the alignment. The letter X in these alignment denotes a lack of conservation at 90% for the particular group at that site whereas a residue means greater than 90% conservation. It should be noted that many alignments had fewer than 10 members and some (with only 1 member) “appear” to be perfectly conserved in the alignment but this is an artefact of the number of group members.

Next we utilised this phylogenetically “collapsed” consensus alignment in Figure S10 to examine each of the key GID1A residues identified in the modelling analysis in the binding pocket, cleft and channel to asses if it was identified as a conserved residue across all plant lineages, conserved across one or more of the four major subclades, conserved across one or more of the subgroupings with these subclades or variable across all 26 divisions. The results of this analysis are presented in supplementary information tables 3, 4, 5.

## Supplemental Information

**Figure S1: RMSD GAI MoRFs during production runs.** RMSD from production runs of GAI MoRF subsets of gDG and aDG systems. Computed from trajectories corrected for PBC via VMD. RMSD frames were averaged over intervals of 300 frames, standard deviation shown in bars.

**Table S1: Logic table for Channel pathway egressions in gG system.** H-bond and Contact residues during Channel pathway egressions gG.

**Table S2: Logic table for Cleft pathway egressions in gDG system.** H-bond and Contact residues during cleft pathway egressions gDG, * interacting GAI residue close to the DELLA motif.

**Figure S2: Initial and final frame for Cleft egression in gG system.** (A) Initial (green/transparent) and final (cyan/solid surface) frames in ribbon display mode from a cleft egression in GID1A with GA_3_. The slight flex or “breathing” motion of helix B, shown in red, in GID1A allows a clear path. (B) Initial (light shading) and final (darker shading) frame of space filling models of the pocket subset (Table 1) showing a cleft egression in GID1A and flexing/breathing of helix B as GA_3_ leaves the cleft.

**Figure S3: RMSF mapped cartoon of GA_3_ DELLA · GID1A egression pathways.** JEDi RMSF maps from example egression paths in GA_3_-DELLA-GID1A system. Perspective of helix B GID1A and DELLA MoRF1.

**Figure S4: RMSF mapped cartoon of GA_3_ GID1A egression pathways.** JEDi RMSF maps of examples egression paths in GA_3_ GID1A system. Perspective of GID1A N-terminal in front.

**Figure S5: RMSD of GA^(x)^ in gG system.**RMSD of GA ligand in gG systems during 20 ns production run.

**Figure S6: Essential motion PCA weights tRAMD for GA_3_-DG complex.** PCA performed at alpha carbon level for GA3-GID1-DELLA tRAMD egressions for examples of the two pathways, first PCA mode showing. Residues numbered linearly from GAI-1 to GID1A-450, see methods for numbering correction. The channel pathway is plotted in red and the cleft pathway in blue. Note that residues 110-450 in this Figure match residues 0 to 340 in SI Figure 7 below.

**Figure S7: Essential motion PCA weights tRAMD for GA_3_-GID1A complex.** PCA performed at alpha carbon level of GA3-GID1A tRAMD egressions for examples of the two pathways. The channel pathway data are plotted in red and the cleft pathway data in blue. Note the y-axis change due to the loss of dynamics in GID1A cleft egression.

**Figure S8: DELLA RMSF comparison between Apo, GA^(a)^ and GA^(i)^ complexes with GID1.** RMSF at alpha carbon level for DELLA subset of gDG (red line with GA^(i)^, black line with GA^(a)^) and DG (apo-pink line) systems.

**Figure S9: RMSF analysis for GID1A in binary and tertiary complexes with DELLA and GA^(a)^.** RMSF at alpha carbon level for GID1A subset of aDG (Apo), gDG,, and gG during 20 ns production run for GA^(a)^ containing systems. gDG and gG systems are averaged with standard deviation shown as bars.

**Table S3: Analysis of contact residues in the binding pocket of GID1 across four main phylogenetic divides.** *1 – Residues underlined indicate H-bond contacts, the others indicate hydrophobic interactions. *2 –denotes residues also identified as important in GA binding by Hao et al. (2013). Frequent (in capitals) and infrequent (in lower case) residue variants are given in brackets using the single letter amino acid code-note the lower case L is underlined to distinguish it from upper case i. C=highly conserved across this clade (in bold indicates a conserved but different residue to that found in the GID1A template), cases of minor variation are given in brackets; S =mainly conserved with conservative substitutions only across this clade.

**Table S4: Analysis of channel contact residues across the main phylogenetic divides of GID1.** *1 – Residues underlined indicate H-bond contacts, the others indicate hydrophobic interactions. *2 –denotes residues also identified as important in GA channel by Hao et al. (2013). Frequent (in capitals) and infrequent (in lower case) residue variants are given in brackets using the single letter amino acid code-note the lower case L is underlined to distinguish it from upper case i. C=highly conserved across this clade (in bold indicates a conserved but different residue to that found in the GID1A template), cases of minor variation are given in brackets; S =mainly conserved with conservative substitutions only across this clade; V*= variable.

**Table S5: Cleft contact residue conservation across 4 main phylogenetic divides of GID1.** *1 – Residues underlined indicate H-bond contacts, the others indicate hydrophobic interactions. Frequent (in capitals) and infrequent (in lower case) residue variants are given in brackets using the single letter amino acid code-note the lower case L is underlined to distinguish it from upper case i. C=highly conserved across this clade (in bold indicates a conserved but different residue to that found in the GID1A template), cases of minor variation are given in brackets; S =mainly conserved with conservative substitutions only across this clade.

**Figure S10: Alignment of 26 GID1 plant and basal plant subclades represented by 90 % conserved consensus.** Geneious alignment of consensus GID1 sequences (created at 90% identity) from 26 sub-clades defined by duplication nodes in plant classification subdivisions in the EnsemblPlants database. All letters denote single letter amino acid code with the exception of X and Z. X denotes residues not conserved in the relevant consensus sequence, Z denotes conserved charged residues at this site. The clade naming convention includes an abbreviated form of the plant family before the GID1 protein name, a major division into GID1A or GID1B in dicot families or indication the clade is a monocot (mon) or more basal (bas) plant family and a final letter (A, B or C) to distinguish multiple subclades with distinct evolutionary histories in the same plant family. The subclades are organised by the most basal subclades at the bottom followed by monocot specific clades, then the GID1B subclades and ending with the GID1A dicot subclades at the top of the alignment to match the tree like phylogeny structure. Moss-Slagenella sp. sequences; Ambt-Amborella trichopoda sequence; Dios-Dioscorea sp. sequences (yam family); Zing-Zingiberales order sequences from Musa sp.; Pani-Panicoideae sub-families within the Poaceae family; Pooi-Pooideae sub-families within the Poaceae family; Oryz - Oryzideae sub-families within the Poaceae family; Cary-Beta sp. sequences in the Caryophyllales order; Lami-Lamiid subclade sp. sequences in the Euasterid clade; Camp-Campanulid subclade sp. sequences in the Euasterid clade; Eric-Ericales order sp. sequences; Vital-Vitales order sequences from Vitis in the SuperRosid clade; Malv-Malvid clade sequence in the Eurosid clade; Fab-Fabid clade sequence in the Eurosid clade.

**Movie S1: PCA motions.** (MP4) The intrinsic dynamics based on Principal Component Analysis associated with PCA mode 1 is shown for the gDG and gG systems when GA3 is bound. The rendering does not include the GA3 molecule. Two movies are shown for the gDG system. The “front” view corresponds to the channel in view, while the “back” view corresponds to the cleft in view. Likewise, two movies are shown for the gG system to show the front and back views. The ribbon rendering has the coloring scheme where blue indicates very little mobility, green indicates low mobility, yellow indicates moderate mobility, orange indicates considerable mobility and red indicates very high mobility.

**Movie S2: Example GA3 egressions.** (MP4) The last part of tRAMD simulations are shown for the case of GA3 dissociating from the channel and cleft pathways in the gDG and gG systems. It is clear there are multiple ways for GA to enter and exit the binding pocket.

**Readme S1: Description of data and script location.**(TXT)

## Acknowledgements

This international collaboration was funded by the New Zealand MBIE (Ministry of Business, Innovation and Employment) Endeavour programme grant C11X1804 to ER. The authors wish to thank editorial contributions from Drs Toshi Foster and Cyril Hamiaux (Plant & Food Research). Donna Gibson (Plant & Food Research) for rendering and conceptualizing art. We wish to acknowledge the use of New Zealand eScience Infrastructure (NeSI) high performance computing facilities in this research. These national facilities are provided by NeSI and funded jointly by NeSI’s collaborator institutions and through the MBIE research infrastructure programme (URL https://www/nesi.org.nz) and made available to other MBIE funded programmes. We also acknowledge the provision of additional computing facilities and resources by the University of North Carolina at Charlotte.

## Author contributions

**Conceptualization**: Charles C. David, Marion Wood, Xiaolin Sun, Donald J. Jacobs, Erik H. A. Rikkerink

**Formal analysis**: John Patterson, Charles C. David, Donald J. Jacobs, Erik H. A. Rikkerink

**Funding acquisition**: Marion Wood, Erik H. A. Rikkerink

**Investigation**: John Patterson, Charles C. David, Marion Wood, Xiaolin Sun, Donald J. Jacobs, Erik H. A. Rikkerink

**Project administration**: Donald J. Jacobs, Erik H. A. Rikkerink

**Supervision**: Donald J. Jacobs, Erik H. A. Rikkerink

**Validation**: Charles C. David, Donald J. Jacobs, Erik H. A. Rikkerink

**Visualization**: John Patterson, Charles C. David, Marion Wood

**Writing – original draft**: John Patterson, Marion Wood, Erik H. A. Rikkerink

**Writing – review & editing**: John Patterson, Charles C. David, Marion Wood, Xiaolin Sun, Donald J. Jacobs, Erik H. A. Rikkerink

